# UPR^ER^–immunity axis acts as physiological food evaluation system that promotes aversion behavior in sensing low-quality food

**DOI:** 10.1101/2023.11.10.566584

**Authors:** Pengfei Liu, Xinyi Liu, Bin Qi

## Abstract

To survive in challenging environments, animals must develop a system to assess food quality and adjust their feeding behavior accordingly. However, the mechanisms that regulate this chronic physiological food evaluation system, which monitors specific nutrients from ingested food and influences food-response behavior, are still not fully understood. Here, we established a low-quality food evaluation assay system and found that heat-killed *E. coli* (HK-*E. coli),* a low sugar food, triggers cellular UPR^ER^ and immune response. This encourages animals to avoid low-quality food. The physiological system for evaluating low-quality food depends on the UPR^ER^ (IRE-1/XBP-1) – Innate immunity (PMK-1/p38 MAPK) axis, particularly its neuronal function, which subsequently regulates feeding behaviors. Moreover, animals can adapt to a low-quality food environment through sugar supplementation, which inhibits the UPR^ER^ –PMK-1 regulated stress response by increasing vitamin C biosynthesis. This study reveals the role of the cellular stress response pathway as physiological food evaluation system for assessing nutritional deficiencies in food, thereby enhancing survival in natural environments.

## Introduction

Food is essential for the survival, growth and fitness of all animals. To adapt to fluctuating environments with a wide range of food sources, animals have developed a food evaluation system. This system enables them to identify nutrient-rich food and avoids low-quality or toxic food, thereby maximizing their survival prospects ^1, 2, 3^. Various sensory neuron evaluation systems in animals has evolved to evaluate food quality through vision ^4, 5^, olfactory ^3, 6, 7, 8, 9, 10, 11,, 12^ and gustatory senses ^13, 14, 15^. Besides these sensory systems that facilitate quick feeding decisions, animals may also initiate cellular stress response programs to detect nutrition/toxin and trigger food response behaviors ^16, 17^. This could be one of physiological food quality evaluation systems that monitor the nutritional status of consumed food. However, the signaling events in cellular stress responses involved in evaluating of specific nutrients and the mechanisms that connect these signaling activities to food behaviors are largely unexplored. More specifically, while cellular stress response through UPR^ER^ ^18^ and PMK-1/p38 MAPK ^19^ dependent immunity in response to pathogens have been extensively studied, the functions of these cellular stress response in sensing and evaluating specific nutrients from food remain unclear.

Vitamin C is an essential micronutrient that cannot be synthesized by humans due to the loss of a key enzyme in the biosynthetic pathway ^20^. Animals obtain vitamin C from their diet and possibly also from gut microbes ^21^. Vitamin C is an important physiological antioxidant and a cofactor for a family of biosynthetic and gene regulatory monooxygenase and dioxygenase enzymes. It is also required for the biosynthesis of collagen, L-carnitine, and certain neurotransmitters ^22, 23^. Vitamin C has been associated with various human diseases including scurvy, immune defect and cardiovascular disease ^20^. Therefore, in animals, having robust food evaluation systems to detect vitamin C levels could significantly impact their survival in the wild. However, the potential involvement of the cellular stress response pathway in this food evaluation system for sensing and assessing vitamin C remains largely unexplored.

In this study, using the low-quality food evaluation assay system we established ^24^, we elucidated the mechanism by which the cellular stress response pathway operates as a physiological food evaluation system. This pathway assesses the deficiency of D-glucose in food and the subsequent vitamin C content in animals through the unfolded protein response (UPR^ER^) – innate immunity (PMK-1/p38 MAPK) axis. This mechanism promotes animals to leave low-quality food and is critical for their survival in nature environments.

## Results

### Low-quality food induces stress response in animals

Our previous studies have shown that Heat-killed *E. coli* (HK-*E. coli*), which lacks certain molecules, is considered a low-quality food that is unable to support animal growth ^24, 25^. Moreover, through metabolic-seq analysis, we identified significant changes in a large numbers of derivatives (Figure 1 – figure supplement 1A, Table S1), including lipids and their derivatives (Figure 1-figure supplement 1B, Table S1), amino acid and their metabolites (Figure 1-figure supplement 1C, Table S1), as well as coenzymes and vitamins (Figure 1-figure supplement 1D, Table S1). Interestingly, we observed a significant decrease in carboxylic acids and their derivatives (Figure 1-figure supplement 1E, Table S1) in *E. coli* after being heat-killed (Figure 1-figure supplement 1F, Table S1). This suggests that HK-*E. coli* is nutritionally deficient for *C. elegans* when compared to normal *E. coli* food.

Next, we conducted two behavior assays to facilitate the analysis of the food evaluation process in animals by seeding L1 animals in assay plates (Figure 1A and 1B). In the avoidance assay, wild-type animals avoided the HK-*E. coli* OP50 (HK-OP50) food (Figure 1A). Interestingly, in the food choice assay, animals initially showed no preference between the two types of food (1-2h), but eventually exhibited a preference for high-quality food (Live *E. coli*) up until the 17h mark (Figure 1B, Figure 1-figure supplement 1G). This suggests that worms depart from the HK-*E. coli* after recognizing it as low-quality food source through ingestion.

**Figure 1.**
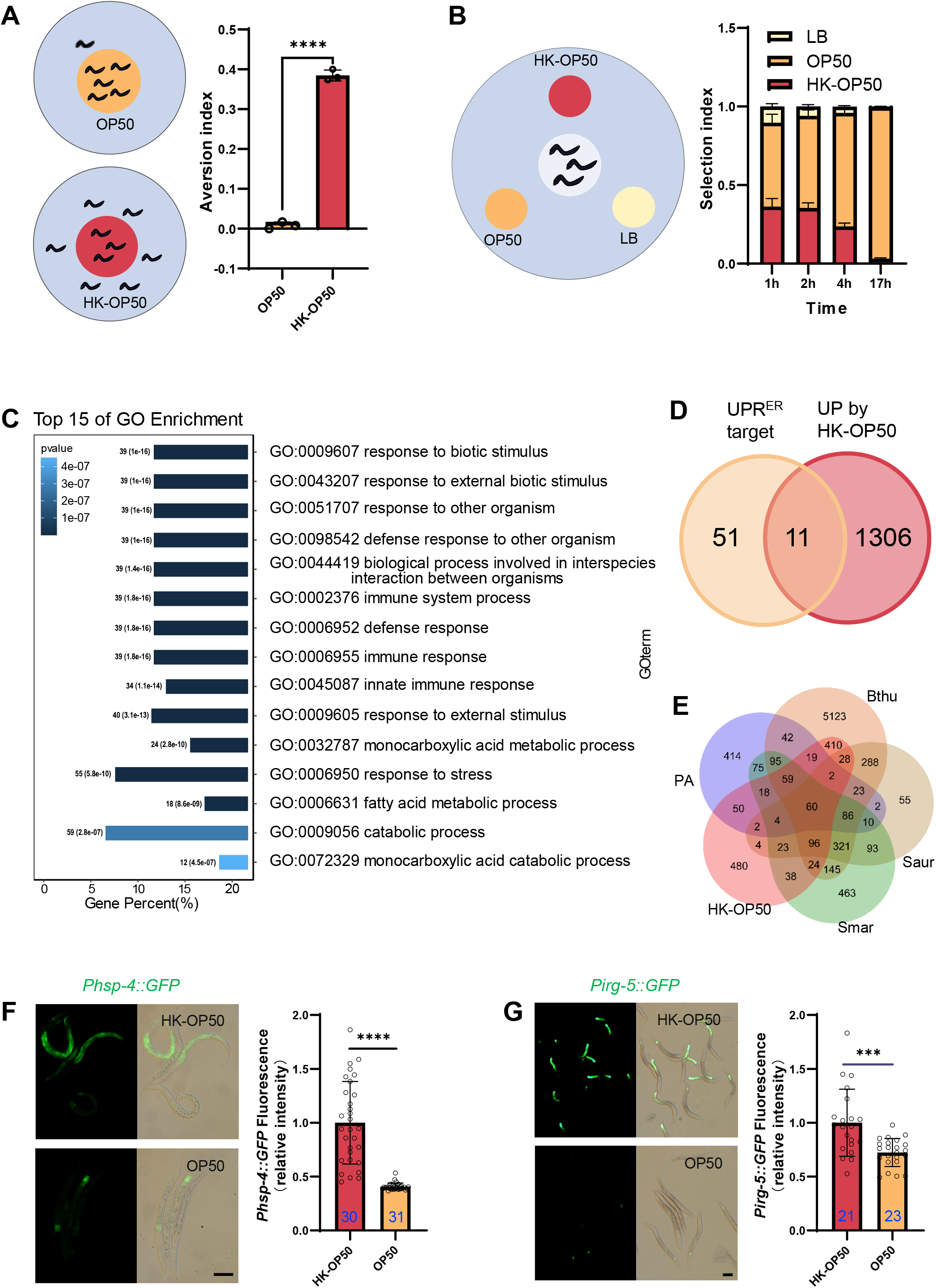
The stress response is induced in animals fed low-quality food, HK-*E. coli*. (A) Schematic drawing and quantitative data of the food aversion assay. Circles indicate the food spot for live (yellow) and HK-OP50 (red) bacteria, respectively. The animals were scored 16-17 hours after L1 worms were placed on the food spot. Data are represented as mean ±SD from three independent experiments, 79-129 animals/assay. (B) Schematic method and quantitative data of the food selection assay. Live (yellow), heat-killed (red) *E. coli* and LB as the buffer for *E. coli* were placed on indicated position. Synchronized L1 worms were place in the center spot. The selection index was calculated at the indicated time. Data are represented as mean ±SD from eight independent experiments, 123-792 animals/assay. (C) GO enrichment analysis of up-regulated genes in animals fed with HK-*E. coli* vs live *E. coli*. (D) Venn diagram showing numbers of UPR^ER^ target genes and up-regulated genes in animals fed HK-*E. coli*, and their overlap. (E) Venn diagram showing numbers of induction genes by four pathogenic bacteria and HK-*E. coli* induced genes, and their overlap. The gene expression data was extracted from published data of animals’ infection with *Pseudomonas aeruginosa* (PA) ^26^, *Bacillus thuringiensis* (Bthu) ^27^, *Staphylococcus aureus* (Saur) ^27^, and *Serratia marcescens* (Smar) ^27^. (F-G) GFP fluorescence images and bar graph showing that *Phsp-4::GFP* (F) and *Pirg-5::GFP* (G) were induced in animals fed with HK-*E. coli.* Blue numbers are the number of worms scored from at least three independent experiments. Data are represented as mean ± SD. For all panels, Scale bar shows on indicated figures, 50 μm. * p<0.05, ** p<0.01, *** p < 0.001, **** p<0.0001, ns: no significant difference. Precise P values are provided in Raw Data.

In order to investigate the pathways in animals that respond to HK-*E. coli*, we performed transcriptomics analysis on worms that were cultured with both HK-*E. coli* and Live *E. coli*. Gene-expression profiling revealed that stress response genes, including those related to biotic stimulus, immune response and response to stress, are up-regulated in animals fed with HK-*E. coli* OP50 (HK-OP50) (Figure 1C, Table S2). Among these up-regulated genes, we identified 11 out of 62 of UPR^ER^ target genes (Figure 1D, Figure 1-figure supplement 1H and Table S2). Additionally, about 50%-80% of up-regulated genes overlap with genes responding pathogenic bacteria ^26, 27^ (Figure 1E, Table S2). Consistent with the results of the RNA sequencing (RNA-seq) analysis, the UPR^ER^ reporter (*Phsp-4::GFP*)^28^ and immunity reporter (*Pirg-5::GFP*)^29^ were strongly induced in intestine (Figure 1F-G) and neurons (Figure 1-figure supplement 2A) by feeding unfavorable food (HK-*E. coli* OP50), suggesting that UPR^ER^ and immune pathways may respond to low-quality food (HK-*E. coli* OP50). As intestinal fluorescence (*Phsp-4::GFP or Pirg-5::GFP)* is easy observation and scoring, the further analyses were done in the intestine.

Moreover, UPR^Mt^ reporter (*Phsp-6::GFP*) ^30^ was weakly induced under HK-*E. coli* feeding condition (Figure 1-figure supplement 2B), and starved worm did not induce UPR^ER^ and immunity (Figure 1-figure supplement 2C-D).

Together, these findings suggest that low-quality food (HK-*E. coli* OP50) triggers a stress response pathway in animals, including UPR^ER^ and innate immune pathway. This implies that animals may assess the quality of food through UPR^ER^ and innate immune pathway.

### Animals evaluate food quality through UPR^ER^-immune-dependent physiological food quality evaluation system

To determine whether the UPR^ER^ and innate immune pathways play a role in evaluating low-quality food, we first examined whether the activation of the UPR^ER^ by HK-*E. coli* was dependent on the known signaling components of the UPR^ER^ branches, including IRE/XBP-1, PERK/ATF-4 and ATF-6 ^31, 32^. We observed no difference in *Phsp-4::GFP* induction with *atf-4* (Figure 2-figure supplement 1A) or *atf-6* (Figure 2-figure supplement 1B) RNAi-mediated knockdown in animals fed with HK-*E. coli*. However, knockdown of *ire-1/xbp-1* or mutation of *xbp-1* reduced GFP fluorescence (Figure 2A, 2B). Among the 11 differentially expressed UPR^ER^ target genes in animals fed with HK-*E. coli* from RNA-seq (Figure 1D), 64% of the genes are IRE-mediated genes (Figure 1 – figure supplement 1H, Table S2). The mRNA level of IRE-1-mediated splicing of *xbp-1*^28^ is also induced in animals fed with HK-*E. coli* OP50 (Figure 2-figure supplement 1C). However, UPR^ER^ is not affect in animals feeding live-*E. coli* by RNAi of *ire-1, xpb-1, atf-4* and *atf-6* (Figure 2-figure supplement 1D). These data suggest that activation of the UPR^ER^ by low-quality food (HK-*E. coli*) depends on the IRE-1/XBP-1.

**Figure 2.**
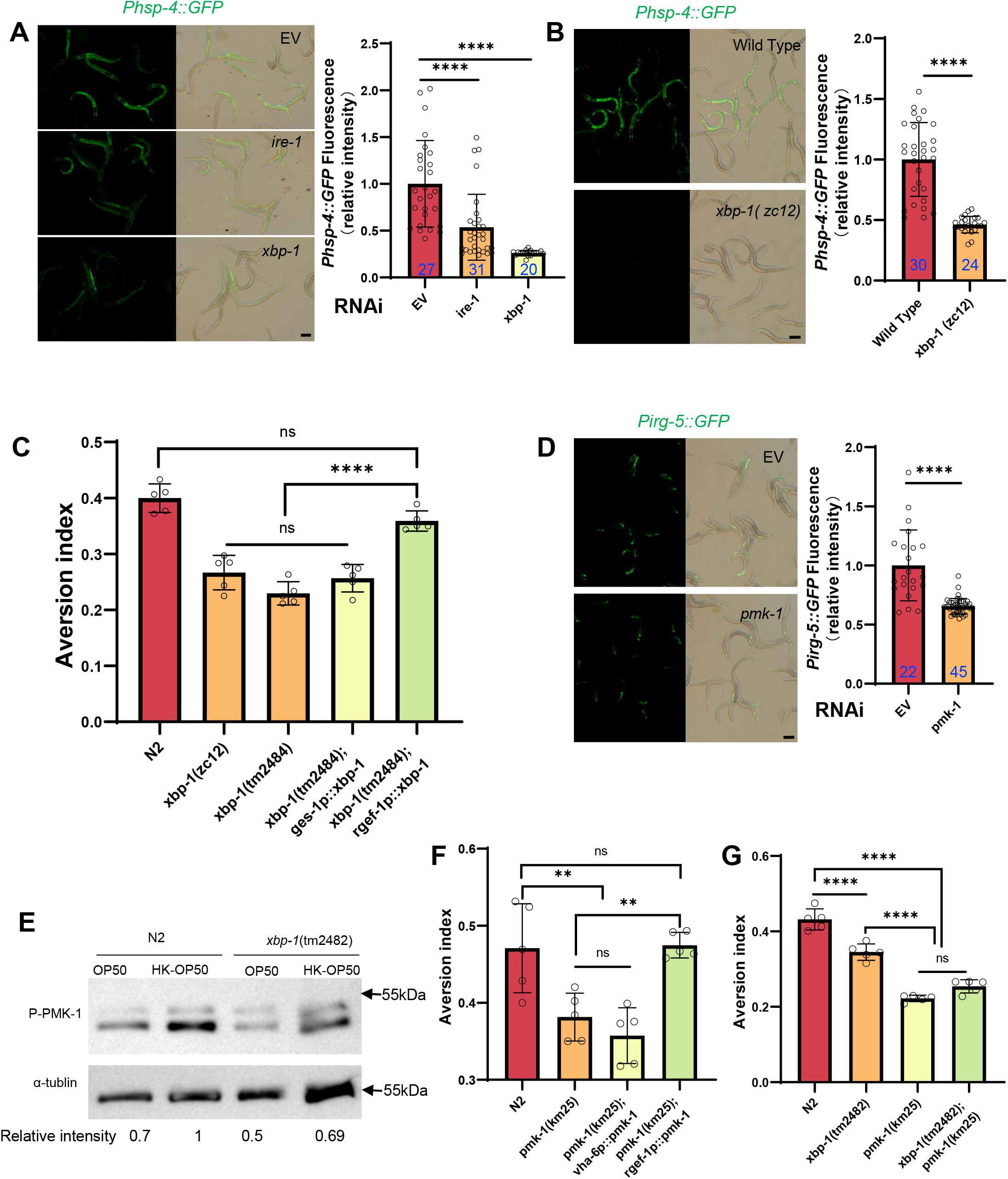
Animals evaluate food quality through UPR^ER^ (*ire-1/xbp-1*) – Innate immunity (*pmk-1* MAPK) axis. (A-B) GFP fluorescence images and bar graph showing that HK-*E. coli* induced *Phsp-4::GFP* was decreased in animals with *ire-1* or *xbp-1* RNAi treatment (A) or *xbp-1* mutation (B). Blue numbers are the number of worms scored from at least three independent experiments. Data are represented as mean ± SD. (C) Food aversion assay showing that *xbp-1* mutation eliminated the discrimination against HK-*E. coli.* However, this effect is rescued by expressing *xbp-1* in neurons rather than intestine. Data are represented as mean ±SD from five independent experiments, 156-763 animals/assay. (D) GFP fluorescence images and bar graph showing that HK-*E. coli* induced *Pirg-5::GFP* was decreased in animals with *pmk-1* RNAi treatment. Blue numbers are the number of worms scored from at least three independent experiments. Data are represented as mean ± SD. (E) Western blot images showing the level of p-PMK-1 in L1 animals (Wild-type N2 and *xbp-1* mutant) fed with OP50 or HK-OP50 for 4 h. The level of p-PMK-1 is induced in animals fed HK-OP50. (F) Food aversion assay showing that *pmk-1* mutation eliminated the discrimination against HK-*E. coli.* However, this effect is rescued by expressing *pmk-1* in neurons rather than intestine. Data are represented as mean ±SD from five independent experiments, 168-492 animals/assay. (G) Food aversion assay in wild-type, *xbp-1*, *pmk-1* and double mutant. Data are represented as mean ±SD from five independent experiments, 259-490 animals/assay. For all panels, Scale bar shows on indicated figures, 50 μm. * p<0.05, ** p<0.01, *** p < 0.001, **** p<0.0001, ns: no significant difference. Precise P values are provided in Raw Data.

To further analyze whether XBP-1-dependent UPR^ER^ activation is critical for animals to leave low-quality food, we tested food avoidance behavior using *xbp-1* mutant. The results show that *xbp-1* mutants had a significantly decreased likelihood of leaving of HK-*E. coli*, which was rescued by expressing *xbp-1* in neuron rather than intestine (Figure 2C). This indicates that XBP-1-dependent UPR^ER^ activation in neuron is critical for animals to specific evaluate low-quality food (HK-*E. coli*).

We then investigated which innate immune pathway is involved in evaluating low-quality food. First, we analyzed HK-*E. coli* induced genes from RNA-seq. Among these up-regulated genes, 82 out of 409 of PMK-1-dependent genes ^33^ were identified (Figure 2-figure supplement 1E, Table S2). Second, we confirmed the induction of several well-known PMK-1 target genes in RNA-seq data^34^ (Figure 2-figure supplement 1F) and reporter analysis (Figure 2-figure supplement 1G-H). Moreover, the induction of *Pirg-5::GFP* was abolished in *pmk-1* knockdown animals fed with HK-*E. coli* (Figure 2D). Third, we found that the phosphorylated PMK-1 (p-PMK-1) level was prominently increased in wild-type N2 animals fed HK-*E. coli* compared to feeding *E. coli* OP50 (Figure 2E). Finally, *pmk-1* mutants had a decreased likelihood of leaving of HK-*E. coli*, which was rescued by expressing *pmk-1* in neurons rather than intestine (Figure 2F). Moreover, *Pirg-5::GFP* is not affect in animals feeding live-*E. coli* by RNAi of *pmk-1* (Figure 2-figure supplement 1I).These data suggest that PMK-1 regulated immune pathway evaluates low-quality food, especially the neuronal PMK-1 has a critical function for food quality response.

### XBP-1 and PMK-1 are in the same pathway for evaluating food quality

Next, we explored the connection between UPR^ER^ (IRE-1/XBP-1) and innate immunity (PMK-1 p38 MAPK) in food quality evaluation. We found that *Pirg-5::GFP* induction (Figure 2-figure supplement 1I, 2A) and PMK-1 activation (Figure 2E) were decreased in animals with *xbp-1* mutation or knockdown when fed with HK-*E. coli*, suggesting that XBP-1 could regulate PMK-1 under this condition. Additionally, *Phsp-4::GFP* induction under HK-*E. coli was* not affected in animals with *pmk-1* RNAi (Figure 2-figure supplement 1D, 2B), indicating that XBP-1-dependent UPR^ER^ activation is not regulated by PMK-1. Finally, we constructed a double mutant of *xbp-1* and *pmk-1* and found that the food avoidance phenotype of the double mutant was similar to the *pmk-1* mutant (Figure 2G), indicating that PMK-1 is downstream of XBP-1 in responding to low-quality food.

We then asked whether UPR^ER^ (IRE-1/XBP-1) – Innate immunity (PMK-1/p38 MAPK) axis is specific to evaluate low-quality food (HK-*E. coli*). We performed behavior assay in N2, *pmk-1* and *xbp-1* mutant animals by feeding normal *E. coli* food, inedible food (*Saprophytic staphylococci)* ^35^ and pathogenic food (*Pseudomonas aeruginosa-PA14)* ^18^. We found that N2, *pmk-1*, and *xbp-1* mutant worms did not exhibit avoidance behavior when presented with normal food (OP50). However, both N2 and *xbp-1* mutant worms were able to escape from inedible food (N2 was predominantly found on the border areas of the bacterial lawn and *xbp-1* mutant worms on border and in), *Saprophytic staphylococci*, whereas *pmk-1* mutant worms did not exhibit this avoidance behavior. Notably, N2 and *xbp-1* mutant worms exhibited even more pronounced avoidance behavior when exposed to *Pseudomonas aeruginosa*, whereas *pmk-1* mutant worms were more susceptible to infection by this pathogen (Figure 2-figure supplement 2C). These findings suggest that the UPR-Immunity pathway plays a crucial role in helping animals avoid low-quality food (HK-*E. coli*) by triggering an avoidance response. In contrast, the Innate immunity pathway, which is mediated by PMK-1/p38 MAPK, appears to play a key role in evaluating unfavorable food sources, such as HK-*E. coli*, *Saprophytic staphylococci*, and *Pseudomonas aeruginosa*, and helping animals avoid these environments.

### Sugar deficiency in HK-*E. coli* food induces stress response and avoidance behavior in animals

We then investigated which nutrients/metabolites are sensed by animals through the XBP-1-PMK-1 axis for food quality evaluation. First, we hypothesized that the nutrition status is improved in *E. coli* mutant (HK-treatment), which could inhibit UPR^ER^ and immune response in animals. We established a system for screening the *E. coli* mutant Keio library (Figure 3 – figure supplement 1A), and identified 20 *E. coli* mutants that did not induce *Phsp-4::GFP* through the UPR^ER^ reporter (*Phsp-4::GFP*) after three rounds of screening (Table S3). From these 20 *E. coli* mutants, we identified 9 *E. coli* mutants that did not induce *Pirg-5::GFP* through the immunity reporter (*Pirg-5::GFP*) screening (Figure 3-figure supplement 1B-C, Table S3). Animals fed HK-*yfbR*, which catalyzes carbohydrate derivative metabolic process^36^, had a decreased ability to leave food (Figure 3A, Table S3), indicating that HK-*yfbR* may be a higher quality food for animals compared to HK-K12.

**Figure 3.**
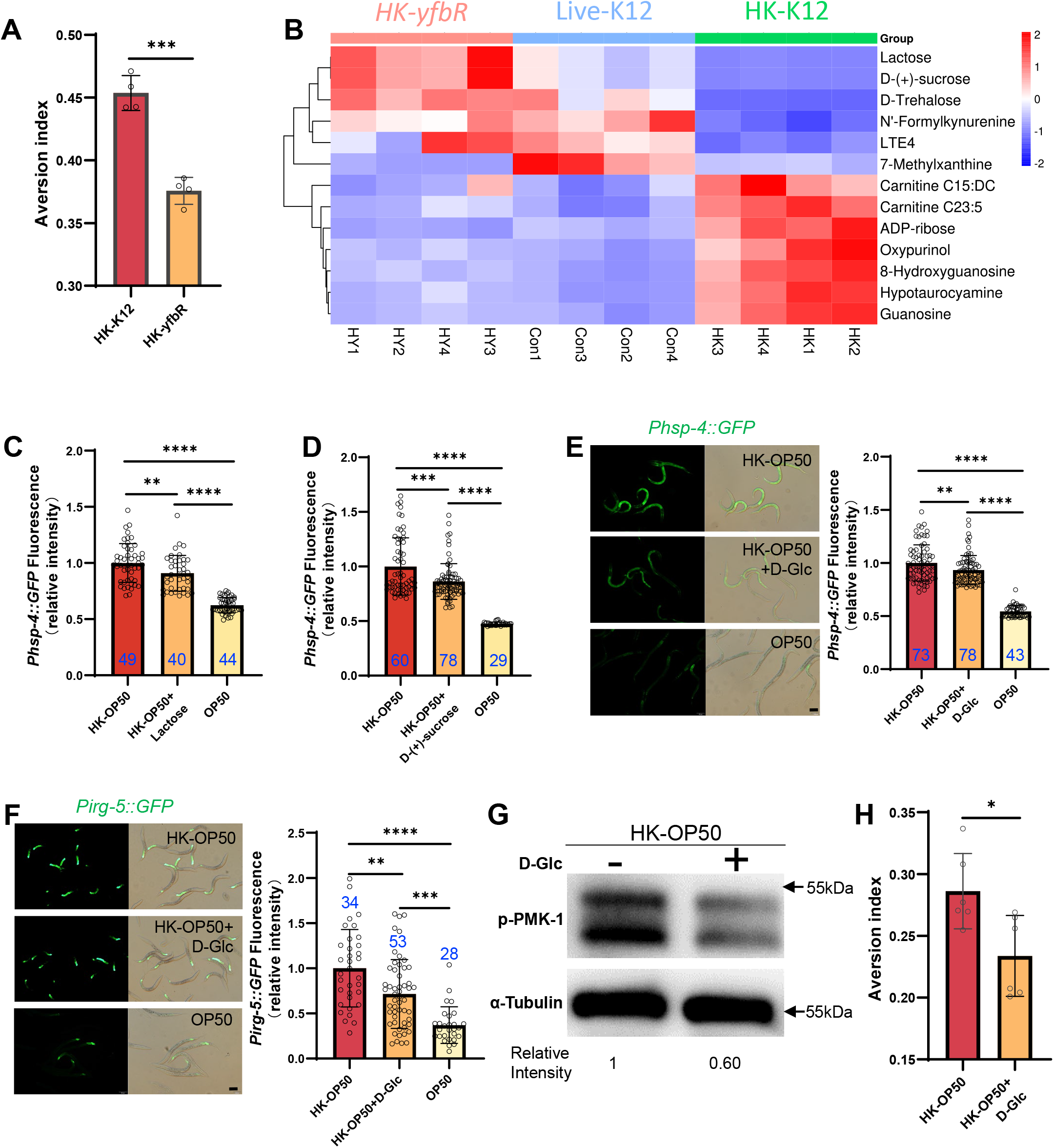
HK-*E. coli* is low sugar food, which induce stress response and avoidance behavior in animals. (A) Food aversion assay showing that wild-type animals eliminated the discrimination against HK-*E. coli* when *yfbR* is mutated in *E. coli*. Data are represented as mean ±SD four independent experiments, 251-490 animals/assay. (B) Heat map showing the thirteen differential metabolites from HK-K12, HK-*yfbR*, and K12 in 4 independent experiments. Color indicates the relative level of each metabolite. (C-D) The bar graph showing that HK-*E. coli* induced *Phsp-4::GFP* was decreased in animals with lactose (C) or D-(+)-sucrose (D) supplementation. Blue numbers are the number of worms scored from at least three independent experiments. Data are represented as mean ± SD. (E-F) GFP fluorescence images and bar graph showing that HK-*E. coli* induced *Phsp-4::GFP* (E) and *Pirg-5::GFP* (F) were decreased in animals with D-(+)-glucose (D-Glc) supplementation. Blue numbers are the number of worms scored from at least three independent experiments. Data are represented as mean ± SD. (G) Western blot images showing the level of p-PMK-1 in L1 animals fed HK-*E. coli* with or without D-(+)-glucose (D-Glc) supplementation for 4 h. The level of p-PMK-1 is decreased in animals fed HK-OP50+D-Glc. (H) Food aversion assay showing that wild-type animals eliminated the discrimination against HK-*E. coli* with D-Glc supplementation. Data are represented as mean ±SD six independent experiments, 190-492 animals/assay. For all panels, Scale bar shows on indicated figures, 50 μm. * p<0.05, ** p<0.01, *** p < 0.001, **** p<0.0001, ns: no significant difference. Precise P values are provided in Raw Data.

Secondly, we performed a metabolomics analysis of different quality food (HK-K12, HK-*yfbR* and Live-K12). We found that the level of 13 metabolites were similar between HK-*yfbR* and Live-K12, but significantly changed in HK-K12 (Figure 3B, Figure 3-figure supplement 2A, and Table S1). We also found that genes involved in glycolysis/gluconeogenesis were up-regulated in animals fed with HK-*E. coli* (Figure 3-figure supplement 2B), suggesting that glycolysis/gluconeogenesis metabolism is disordered in animals fed with HK-*E. coli,* which may result from changes in sugar/carbohydrate intake. The carbohydrates (D-trehalose, lactose, and D-(+)-sucrose) were also decreased in HK-*E. coli* (Figure 3B), suggesting that carbohydrate deficiency may induce stress response and avoidance behavior in animals feeding HK-*E. coli*.

Thirdly, to determine which carbohydrate inhibits stress response in animals, we supplemented each metabolite to HK-*E. coli* and found that only Lactose, and D-(+)-sucrose inhibited HK-*E. coli* induced UPR^ER^ (Figure 3C and 3D, Figure 3-figure supplement 2C). Moreover, we found from our metabolomic data that the sugar level, including lactose, and D-(+)-sucrose, and D-(+)-glucose, was also decreased in HK-*E. coli* (Figure 3B, Table S1). Since lactose and D-(+)-sucrose are hydrolyzed to produce glucose^37, 38^, we wondered whether glucose also inhibits the stress response in animals. We found that D-(+)-glucose supplementation also inhibited HK-*E. coli* induced UPR^ER^ (Figure 3E), immune response (Figure 3F, 3G and Figure 3-figure supplement 2D) and avoidance (Figure 3H). Moreover, sugar supplementation did not affect UPR^ER^ and immunity in normal food (OP50) or starved condition (NGM) (Figure 3 – figure supplement 2E-F). While sugar effectively inhibits the HK-*E. coli*-induced UPR^ER^ and immune response, it does not fully suppress it to the extent observed with live-*E. coli* (Figure 3C-F). This implies that additional nutrients present in live-*E. coli* might also contribute to the inhibition of UPR^ER^ and immune response.

Previous studies have shown that heat-killed *E. coli* (HK-*E. coli*) is a low-quality food source that cannot support the growth of *C. elegans* larvae^24, 25^, whereas supplementation with vitamin B2 (VB2) can restore animal growth^24^. Here, we found that sugar deficiency in HK-*E. coli* induces the UPR^ER^-immune response and avoidance behavior in *C. elegans*. Given this, we investigated whether sugar supplementation could promote animal growth when fed HK-*E. coli.* To our surprise, supplementing HK-*E. coli* with carbohydrates (D-Glc, D-GlcA) did not support animal development (Figure 3-figure supplement 2G), suggesting that carbohydrates are not essential for supporting animal growth on this food source. However, we did find that carbohydrates are critical for inhibiting the UPR^ER^-immune response induced by sugar deficiency in HK-*E. coli*.

Together, these findings suggest that HK-*E. coli* induces a stress response and avoidance behavior in animals, which can be inhibited by D-(+)-glucose supplementation. This implies that animals may evaluate the sugar deficiency from HK-*E. coli* through the activation of UPR^ER^ and immune responses.

### Animals could overcome a low-quality food environment by sugar supplementation through vitamin C biosynthesis

We discovered that D-(+)-glucose supplementation inhibited HK-*E. coli* induced UPR^ER^ (Figure 3E), immune response (Figure 3F, 3G and Figure 3-figure supplement 2D) and avoidance (Figure 3H). Simultaneously, vitamin C (VC), which is synthesized by glucuronate pathway using D-glucose ^39, 40^ (Figure 4A), was found to contribute to neuroprotective^41, 42^, immune defense^43, 44^, and inhibits inflammatory and ER stress^45, 46^. This led us to question whether the vitamin C biosynthesis pathway is involved in evaluating low-quality food by using D-glucose.

**Figure 4.**
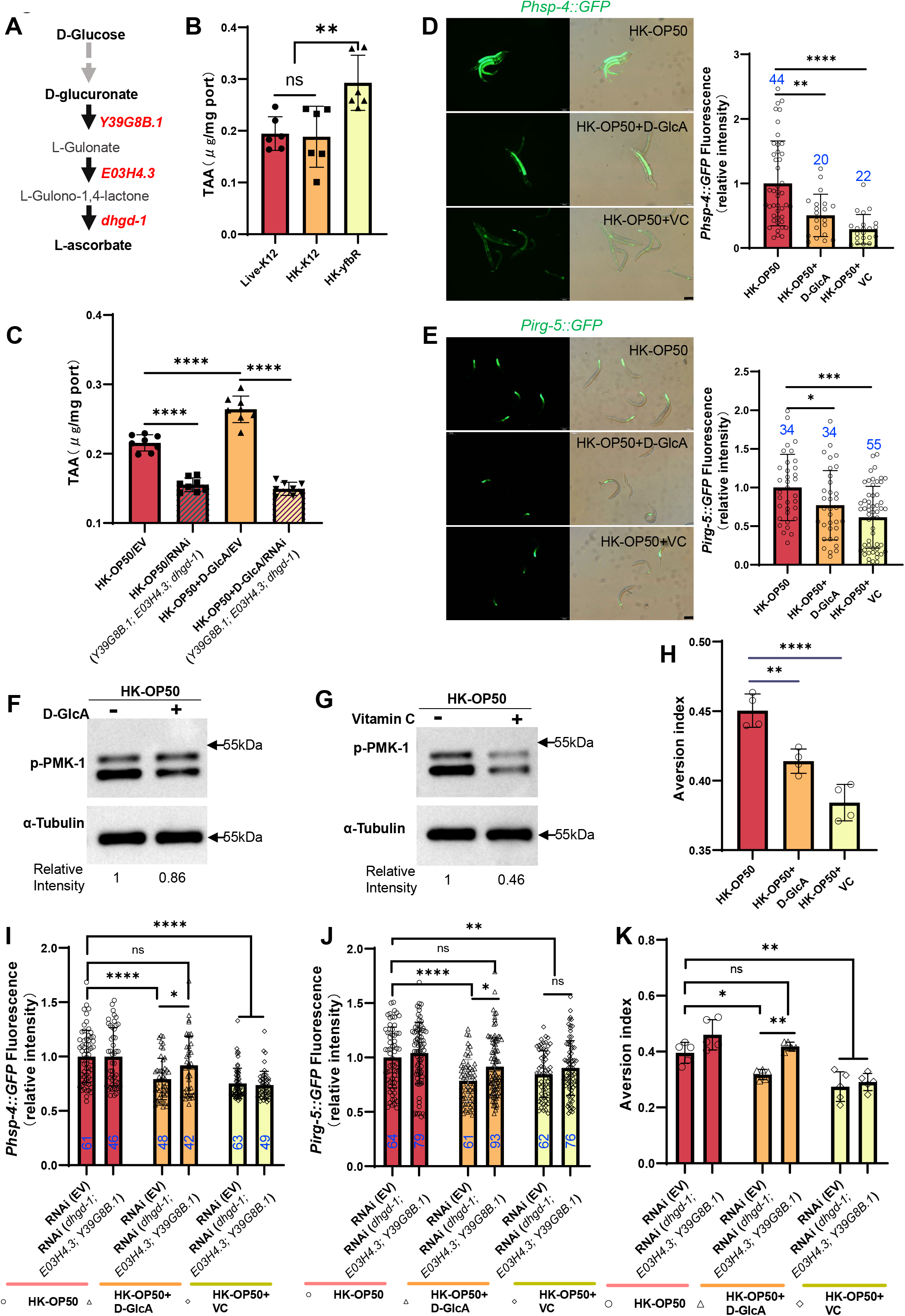
Vitamin C biosynthesis pathway is critically involved in evaluating sugar in the food. (A) Cartoon illustration of a simplified, Vitamin C biosynthesis pathway in *C. elegans*. The relevant coding genes of enzymes was labeled with red. (B) The level of total L-ascorbic acid (TAA), also known as vitamin C, in animals fed with Live-K12, HK-K12, or HK-*yfbR*. Data are represented as mean ±SD from six independent experiments. (C) The level of total L-ascorbic acid (TAA) in animals (control or knockdown of Vitamin C biosynthesis genes) fed with HK-*E. coli* with or without D-glucuronate (D-GlcA) supplementation. Data are represented as mean ±SD from eight independent experiments. (D-E) GFP fluorescence images and bar graph showing that HK-*E. coli* induced *Phsp-4::GFP* (D) and *Pirg-5::GFP* (E) were decreased in animals with D-GlcA or Vitamin C supplementation. Blue numbers are the number of worms scored from at least three independent experiments. Data are represented as mean ± SD. (F-G) Western blot images showing the level of p-PMK-1 in L1 animals fed with HK-*E. coli* with D-GlcA or Vitamin C supplementation for 4 h. The level of p-PMK-1 is decreased in animals with D-GlcA (F) or Vitamin C (G) supplementation. (H) Food aversion assay showing that wild-type animals eliminated the discrimination against HK-*E. coli* with D-GlcA or Vitamin C supplementation. Data are represented as mean ±SD from four independent experiments, 153-292 animals/assay. (I-K) The bar graph showing that suppression of HK-*E. coli* induced *Phsp-4::GFP* (I), *Pirg-5::GFP* (J) and food avoidance (K) by D-GlcA supplementation was abolished in animals with RNAi of VC biosynthesis genes, which was not affect by Vitamin C supplementation. Blue numbers are the number of worms scored from at least three independent experiments and Data are represented as mean ±SD(I-J). Data are represented as mean ±SD from five independent experiments, 252-537 animals/assay (K). For all panels, Scale bar shows on indicated figures, 50 μm. * p<0.05, ** p<0.01, *** p < 0.001, **** p<0.0001, ns: no significant difference. Precise P values are provided in Raw Data.

Firstly, we observed an increase in the vitamin C level in *C. elegans* when fed with HK-*yfbR* (Figure 4B), a high carbohydrate food compared to HK-*E. coli* (Figure 3B and Figure 1-figure supplement 1E). However, the VC level in bacteria is the same (Figure 4-figure supplement 1A). The VC level also increased when D-glucose (Figure 4-figure supplement 1B) or D-glucuronate (D-GlcA) was added to HK-*E. coli* (Figure 4C), which was abolished by knocking down VC biosynthesis genes (Figure 4C and Figure 4-figure supplement 1B). This suggests that addition of sugar supplementation promotes VC synthesis in animals fed with HK*-E. coli*.

Secondly, we hypothesized that animals could overcome a low-quality food (HK-*E. coli*) environment by inhibiting the stress response through increasing vitamin C biosynthesis. We found that VC or D-glucuronate (D-GlcA) supplementation inhibits HK-*E. coli* induced UPR^ER^ (Figure 4D), immune response including *irg-5/sysm-1* reporter expression (Figure 4E and Figure 4 – figure supplement 1C) and p-PMK-1 (Figure 4F and 4G), as well as food avoidance (Figure 4H).

Finally, we asked whether inhibition of stress response and avoidance by sugar supplementation depends on the vitamin C biosynthesis pathway. We found that suppression of HK-*E. coli* induced UPR^ER^ (Figure 4I), immune response (Figure 4J and Figure 4-figure supplement 1D) and food avoidance (Figure 4K) by D-GlcA/sugar supplementation was abolished in animals with RNAi of VC biosynthesis genes. Food selection behavior assays showed that D-GlcA/sugar supplementation increased the preference for heat-killed bacteria, which was also suppressed by knocking down VC biosynthesis genes (Figure 4-figure supplement 1E). However, VC supplementation still suppressed the UPR^ER^ (Figure 4I), immune response (Figure 4J and Figure 4-figure supplement 1D) and food avoidance (Figure 4K), and increased the food preference (Figure 4 – figure supplement 1E) in animals with or without RNAi of VC biosynthesis genes. This suggests that VC, as the final metabolite synthesized from D-glucose, is critical for evaluating low-quality food response in animals.

Together, these data indicate that the vitamin C biosynthesis pathway is critical for evaluating whether food is of higher quality and can be eaten by animals. It also suggests that animals could improve their VC levels to adapt to bad food environment.

### Animals evaluate sugar and vitamin C through neuronal XBP-1 and PMK-1

As D-GlcA/sugar and VC supplementation suppressed HK-*E. coli* induced UPR^ER^, immune response and food avoidance behavior, we investigated whether animals evaluate sugar and VC through XBP and PMK-1 dependent pathways. We performed a food selection behavior assay by adding D-Glc, D-GlcA or VC to the NGM, *E. coli* and HK-*E. coli* (Figure 5A). The food selection behavior assays revealed that supplementation with D-Glc, D-GlcA, or VC inhibits the animals’ choice of sugar or VC on *E. coli*-OP50 feeding conditions (Figure 5-figure supplement 1A). This suggests that supplementation with D-Glc, D-GlcA, or VC may alter the metabolites of live bacteria, leading to avoidance by the animals. There was no preference observed on NGM plate (no food condition) supplementation with D-Glc and VC (Figure 5-figure supplement 1B), indicating that the intake of sugar or VC alone does not influence animal preference. However, alone D-GlcA could influence worm physiology which induces preference change (Figure 5-figure supplement 1B). Interestingly, D-Glc and D-GlcA (Figure 5B and 5C) or VC (Figure 5D) supplementation increased the preference for heat-killed bacteria, which was suppressed in *xbp-1* or *pmk-1* mutant animals. However, this preference was also rescued in *xbp-1* or *pmk-1* mutant animals by expressing XBP-1 or PMK-1 in neurons rather than intestine (Figure 5B-D), indicating that neuronal XBP-1 and PMK-1 are critical for physiological food elevation system for monitoring the level of sugar and VC under low-quality food condition.

**Figure 5.**
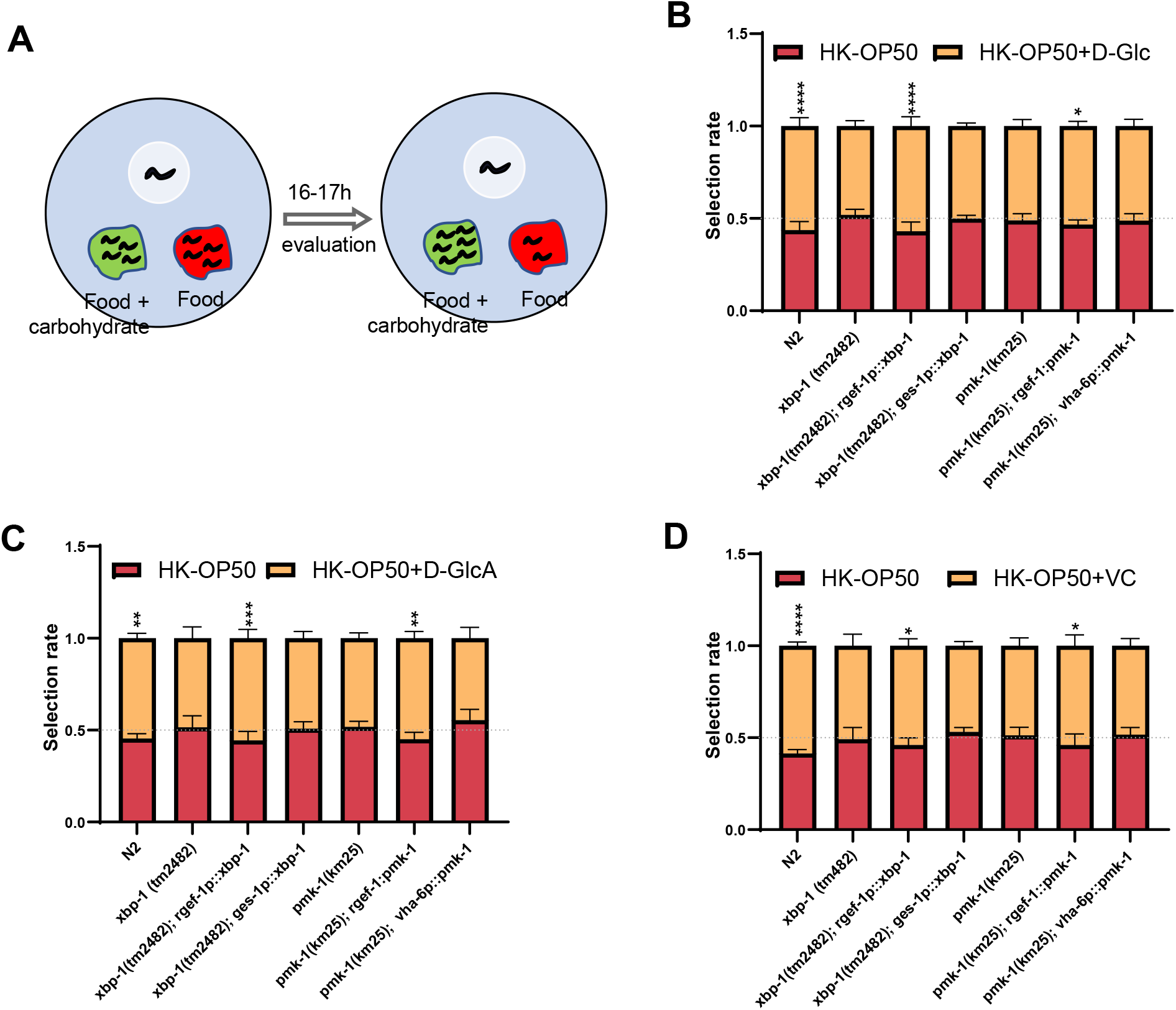
Animals evaluate sugar and vitamin C through neuronal XBP-1 and PMK-1. (A) Schematic method of the food selection assay. Food (red) and food with carbohydrate (D-Glc, D-GlcA, or VC) supplementation (green) was placed on indicated position. Synchronized L1 worms were then place in plate. After 16-17hs, the selection index was calculated. (B-D) Food selection assay showing that *xbp-1* or *pmk-1* mutation eliminated the preference of HK-*E. coli* with D-Glc (B), D-GlcA (C) or Vitamin C (D) supplementation, which was rescued in *xbp-1* or *pmk-1* mutant animals by expressing XBP-1 or PMK-1 in neurons rather than intestine. Data are represented as mean ±SD from five independent experiments, 68-647 animals/assay (B). Data are represented as mean ±SD from six independent experiments, 83-701 animals/assay (C). Data are represented as mean ±SD from six independent experiments, 67-1035 animals/assay (D). For all panels, Scale bar shows on indicated figures, 50 μm. * p<0.05, ** p<0.01, *** p < 0.001, **** p<0.0001, ns: no significant difference. Precise P values are provided in Raw Data.

## Discussion

To better survive, animals must evolve a system to recognize and evaluate the quality of their food. This includes the sensory neuron evaluation system for immediate response and feeding decision ^2^, as well as physiological food evaluation system for chronic response to ingested food. In our previous study, we discovered that the TORC1-ELT-2 pathway, acting as master regulators in intestine, evaluates vitamin B2 deficiency in low-quality food (HK-*E. coli)* and regulates gut digestive activity to impact animal’s food behavior ^24^. To further identified the mechanism by which animals evaluate low-quality food (HK-*E. coli*), we performed metabolomics and transcriptomics analyses to identify specific nutrition deficiencies in low-quality food and the cellular response pathways that are involved in food evaluation pathway. This study identified a physiological food evaluation mechanism by which animals recognize food quality through UPR^ER^ (IRE-1/XBP-1) – Innate immunity (PMK-1/p38 MAPK) regulated cellular stress response program in neurons that dictates food avoidance and selection behaviors (Figure 6).

**Figure 6.**
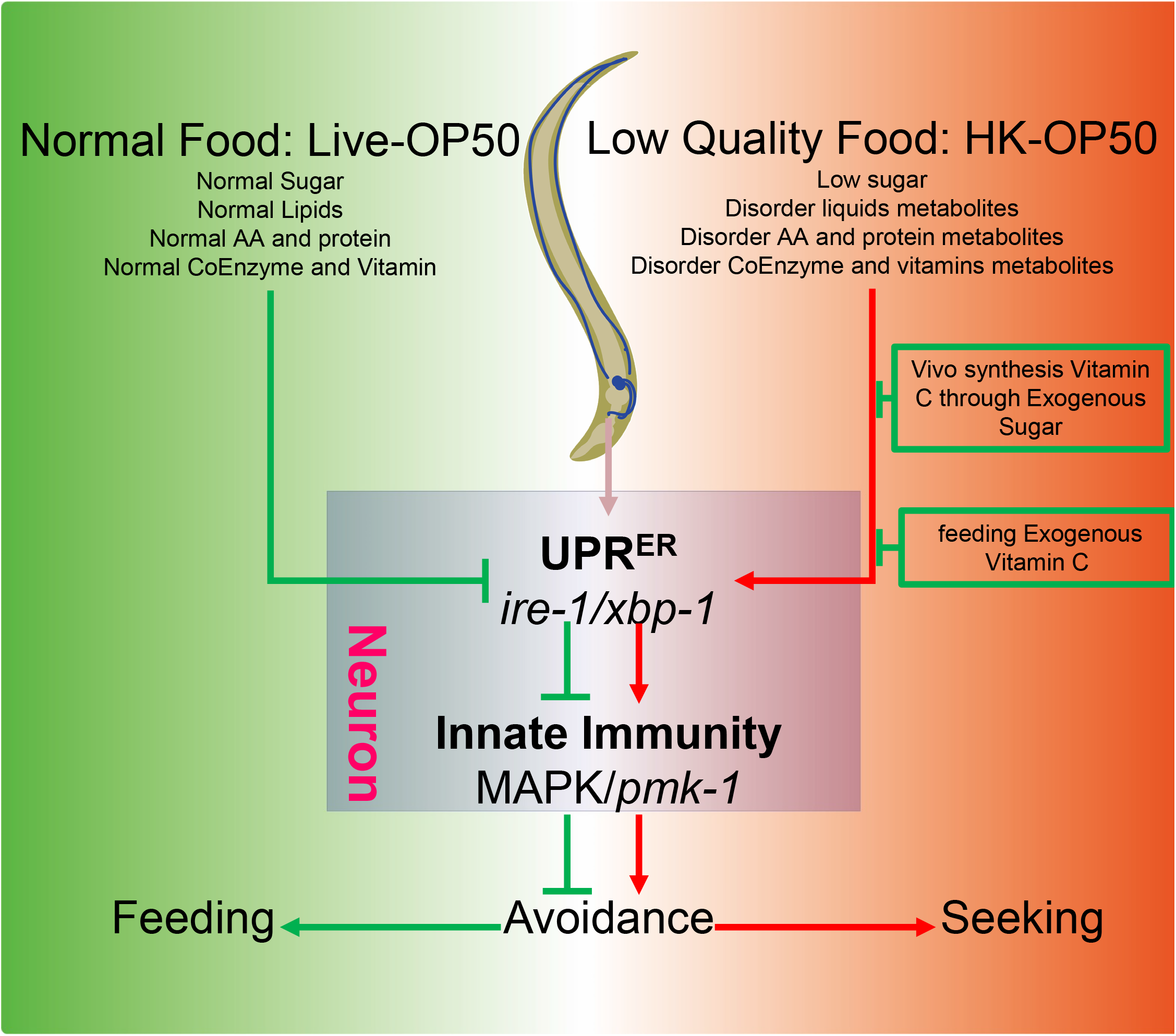
Schematic model of physiological food evaluation system in evaluating/sensing sugar and vitamin C through UPR^ER^ (IRE-1/XBP-1) – Innate immunity (PMK-1/p38 MAPK) axis. Vitamin C level is low in animals fed low sugar food, HK-*E. coli*. Sugar and Vitamin C deficiency activate cellular UPR^ER^ and immune response, which promote animals to leave low-quality food and seek better food for survival. This cellular stress regulated physiological food evaluation system depends UPR^ER^ (IRE-1/XBP-1) – Innate immunity (PMK-1/p38 MAPK) axis in neuron.

One of the cellular stress response pathways, the Unfolded Protein Response (UPR^ER^), is activated by various stresses, including infection and nutrition deficiency, which disrupt the homeostasis of the endoplasmic reticulum (ER)^31^. The activation of UPR^ER^, specifically in the nervous system, has been shown to promote changes in feeding and foraging behavior^47^. The p38 PMK-1 pathway is also crucial for regulating the expression of secreted innate immune effectors and is essential for survival during infection^48^. Therefore, these two pathways play a critical role in ensuring animals’ survival in changing environments. However, it is still unclear whether UPR^ER^ and innate immunity evaluate food quality under physiological conditions. Our study provides evidence that low-quality food (HK-*E. coli*) activates both UPR^ER^ and p-PMK-1, leading to animals leaving the low-quality food. Previously finding have shown that inhibition of ER function promotes *C. elegans* avoid to the toxic food, which employs the MLK-1/MEK-1/KGB-1 pathway^49^. Notably, HK-*E. coli* induced avoidance behavior is independent of the KGB-1 pathway (Figure 5-figure supplement 1C). Additionally, our study reveals that neuronal UPR^ER^ and PMK-1 are essential for evaluating low-quality food, suggesting that the nervous system plays a critical role in assessing food quality.

Previous studies have shown that XBP1 deficiency in intestinal epithelial cells leads to IRE1a hyperactivation and increased JNK phosphorylation in the epithelial compartment in vivo^50^. The IRE1-XBP1 axis has been identified as a critical protective branch of the Unfolded Protein Response (UPR) induced secondary to an innate immune response in the presence of *P. aeruginosa* ^18, 51^. The p38 MAPK has also been shown to directly act on the phosphorylation of IRE-1 to promote the stress response^52, 53^. Interestingly, IRE-1 has been found to confer cold resistance independently of XBP-1 by activating JNK-1 MAPK^49^. In contrast, our study reveals a new mechanism where the UPR^ER^ (IRE-1/XBP-1) positively regulates Innate Immunity (PMK-1/p38 MAPK) under HK-*E. coli* food conditions, establishing a novel physiological food evaluation system that activates the cellular stress response program.

A previous study has shown that activating innate immunity (PMK-1 MAPK) leads to a reduction in translation^54^. Our own previous research has also demonstrated that PMK-1 activation causes a shutdown of food digestion in animals^35^, likely to reduce protein translation and cellular metabolism. To investigate this further, we measured the translation level of animals fed with HK-*E. coli* and found that total translation ability is significantly reduced in these animals (Figure 5-figure supplement 1D). This finding suggests that activating innate immunity (PMK-1 MAPK) may serve as a mechanism to slow down translation progress, thereby alleviating the pressure on the unfolded protein response (UPR) and preventing excessive UPR^ER^ activation.

Vitamin C (VC) is an important physiological antioxidant and a cofactor for a family of biosynthetic and gene regulatory monooxygenase and dioxygenase enzymes. It is also required for the biosynthesis of collagen, L-carnitine, and certain neurotransmitters ^22, 23^. Meanwhile, VC helps animals to protect neuron^41, 42^, defend excessive immune ^43, 44^, and inhibit inflammatory and ER stress^45, 46^ in order to better survive. The synthesis of vitamin C (VC) occurs through the glucuronate pathway, utilizing D-glucose as a precursor ^39, 40^ (Figure 4A). This led us to investigate whether the vitamin C biosynthesis pathway is involved in evaluating low-quality food by using D-glucose. In this study, we found that animals feeding live *E. coli*, which should produce more VC, exhibit higher glucose levels. However, our results show that animals maintain similar VC levels when fed ideal food (live *E. coli*) compared to low-quality food (HK-*E. coli*) (Figure 4B), suggesting that animals do not stimulate VC biosynthesis under favorable food conditions. In contrast, when animals are fed low-quality food (HK-OP50), we found that supplementing D-GlcA in HK-*E. coli* or *E. coli*-*yfbR* mutation can improve VC levels and inhibit UPR^ER^-immunity (Figure 4C). These data indicate that glucose boosts the animal’s ability to adapt to unfavorable food environments by increasing VC levels, but not in favorable food conditions.

Unlike the sensory neuron evaluation system, which permits rapid feeding decisions through smell and taste, the cellular stress response as physiological food evaluation system describe here requires a slow and multi-step signal transduction process after the ingestion of food. The disruption of cellular homeostasis by ingested of low-quality or toxic food can activate stress response mechanisms that both increase the cellular ability to withstand and adapt to this disruption of homeostasis and promote behavioral strategies to avoid these conditions and lessen their impact on the organism. These cellular stress response mechanisms include heat shock response, unfolded protein response, oxidative stress response ^55^. Therefore, this slow physiological food evaluation system is an evolutionary adaptation mechanism for detecting nutrition deficiencies in food that was not detected by quick sensory nervous system.

One limitation of our study is the lack of explanation for why HK-*E. coli* activates UPR^ER^ and immunity. We hypothesized that when heat-killed, HK-*E. coli* may lack or contain altered levels of certain metabolites that either activate or inhibit UPR^ER^ and immunity, respectively. Additionally, we speculated that *E. coli* mutants killed by heat may lack metabolites that activate UPR^ER^ and immunity, or conversely, have increased levels of metabolites that inhibit these pathways. Fortunately, our investigation led to the discovery of the *E. coli* mutant *yfbR*, which inhibits UPR^ER^ and immunity by increasing carbohydrates that aid in resisting these stress pathways. Moving forward, we intend to further explore the intricate relationship between HK-*E. coli* and UPR^ER^-immunity. This will be a key focus of our future research efforts.

Collectively, this study uncovers the unexpected function of UPR^ER^ (IRE-1/XBP-1) – Innate immunity (PMK-1/p38 MAPK) as a physiological food evaluation system for evaluating and sensing food quality in animals. It also highlights the utility of the HK-*E. coli* (low-quality food) – *C. elegans* interaction as a means to dissect the mechanism of food evaluation system in assessing food. Most importantly, it reveals that animals are capable of altering their nutrient (Vitamin C) levels through in vivo synthesis or food intake to adapt to a poor food environment when better food choices are not available.

### Author Contributions

P. L designed, performed experiments, analyzed data. X. L constructed all transgenic animals. B.Q. designed research, supervised this study, and wrote the paper with inputs from P. L.

## Acknowledgments

We thank the Caenorhabditis Genetics Center (CGC) (funded by NIH P40OD010440) for strains; Dr. Zhao Shan for suggestions. This work was supported by the Ministry of Science and Technology of the People’s Republic of China (2019YFA0802100, 2019YFA0803100), the National Natural Science Foundation of China (32170794), Yunnan Applied Basic Research Projects (202302AP370005, 202201AT070196, K264202230211, 202001AV070011), the Yunnan University Startup Program.

## Declaration of interests

The authors declare no competing interests.

## Supplemental Figure legends

**Figure 1 – figure supplement 1.** Food selection assay of animals fed HK-*E. coli* or *E. coli.* Relative to Figure 1. (A-E) Metabolomics analysis of different quality food (HK-K12, HK-*yfbR* and Live-K12). Cluster analysis of all metabolites (A), lipids and their derivatives (B), amino acids and their derivatives (C), coenzymes and vitamins (D), and carbohydrates and their derivatives (E) from Live-K12, HK-K12, and HK-*yfbR*. Color indicates the relative level of each metabolite. HK-K12: heat-killed *E. coli* wild-type K12; HK-*yfbR:* heat-killed *E. coli mutant yfbR;* K12: live *E. coli* wild-type K12. z-score for standardizing data, complete for bi-clustering algorithm, and Euclidean for distance method. (F) Principal component analysis to test the repeatability of the metabolic experiment. HK-K12: heat-killed *E. coli* wild-type K12; HK-*yfbR*: heat-killed *E. coli* mutant *yfbR*; K12: live *E. coli* wild-type K12. LB: LB buffer for culturing *E. coli*. (G) The number of worms in each position calculated at the indicated time, indicating that animals initially select both foods (1-2h), but eventually favor high-quality food (Live E. coli) until 17h. (H) GO enrichment analysis of UPR^ER^ dependent IRE-1 branch. For all panels, * p<0.05, ** p<0.01, *** p < 0.001, **** p<0.0001, ns: no significant difference. Precise P values are provided in Raw Data.

**Figure 1 – figure supplement 2.** Stress response of animals fed HK-*E. coli* or *E. coli.* Relative to Figure 1. (A) UPR^ER^ reporter (*Phsp-4::GFP*) and immunity reporter (*Pirg-5::GFP*) were induced in the neuron and intestine of animals fed with HK-*E. coli.* Neurons are highlighted with a red arrow. (B) UPR^mit^ reporter (*Phsp-6::GFP*) was weakly induced in animals fed with HK-*E. coli*. (C) UPR^ER^ reporter (*Phsp-4::GFP*) expression in animals under normal food (OP50), low quality food (HK-OP50), and starved (M9: hatching L1 worm in M9, NGM) condition; L1 animals were cultured in OP50, HK-OP50 or starved NGM for 20h. (D) Immunity reporter (*Pirg-5::GFP*) expression in animals under normal food (OP50), low quality food (HK-OP50), and starved (M9: hatching L1 worm in M9, NGM). L1 animals were cultured in OP50, HK-OP50 or starved NGM for 20h. For all panels, Scale bar shows on indicated figures, 50 μm.

**Figure 2 – figure supplement 1.** UPR^ER^ and innate immunity pathway in animals are critical for evaluating HK-*E. coli*. Relative to Figure 2. (A-B) GFP fluorescence images and bar graph showing that HK-*E. coli* induced *Phsp-4::GFP* was not affected in animals with *atf-4* (A) or *atf-6* (B) RNAi treatment. Blue numbers are the number of worms scored from at least three independent experiments. Data are represented as mean ± SD. (C) qPCR showing that IRE-1-mediated splicing of *xbp-1* mRNA is induced in animals fed with HK*-E. coli*. (D) UPR^ER^ reporter (*Phsp-4::GFP*) expression in animals with candidate RNAi feeding OP50 or HK-OP50. (E) Venn diagram showing the numbers of PMK-1 dependent genes ^30^ and up-regulated genes in animals fed HK-*E. coli,* and their overlap. (F) The expression of PMK-1 dependent genes which was extracted from RNA-seq data from animals fed with HK*-E. coli*. The data from average of three independent experiments. (G-H) GFP fluorescence images and bar graph showing that *Psysm-1::GFP* (G) and *Pirg-1::GFP* (H) were induced in animals fed HK*-E. coli.* Blue numbers are the number of worms scored from at least three independent experiments. Data are represented as mean ± SD. (I) Immunity reporter (*Pirg-5::GFP*) expression in animals with candidate RNAi feeding OP50 or HK-OP50. For all panels, Scale bar shows on indicated figures, 50 μm. * p<0.05, ** p<0.01, *** p < 0.001, **** p<0.0001, ns: no significant difference. Precise P values are provided in Raw Data.

**Figure 2 – figure supplement 2.** UPR^ER^ positively regulates innate immunity pathway in animals. Relative to Figure 2. (A) GFP fluorescence images and bar graph showing that HK-*E. coli* induced *Pirg-5::GFP* was decreased in animals with *ire-1* or *xbp-1* RNAi treatment. Blue numbers are the number of worms scored from at least three independent experiments. Data are represented as mean ± SD. (B) GFP fluorescence images and bar graph showing that *Phsp-4::GFP* was not affected in animals with *pmk-1* RNAi treatment. Blue numbers are the number of worms scored from at least three independent experiments. Data are represented as mean ± SD. (C) Images and bar graph showing that the avoidance behavior of N2, *xbp-1* mutant, and *pmk-1* mutant in response to different food source (*Saprophytic staphylococci-*SS, *Pseudomonas aeruginosa*-PA14, or OP50). “Broder” indicates regions with thicker SS boundaries; “In” denotes areas inside SS; “Out” refers to areas without SS. The scale is as follows: 2 for animals capable of escaping from PA14 and feeding on PA14 located with edges; 1 for animals capable of escaping from PA14 but not feeding on PA14 located with edges; 0 for animals unable to escape from PA14 and being killed by PA14. Data are represented as mean ±SD from five independent experiments, 359-670 animals/assay. For all panels, Scale bar shows on indicated figures, 50 μm. * p<0.05, ** p<0.01, *** p < 0.001, **** p<0.0001, ns: no significant difference. Precise P values are provided in Raw Data.

**Figure 3 – figure supplement 1.** *E. coli* Keio mutant screening. Relative to Figure 3. (A) Flow chart of strategy for *E. coli* Keio mutant screening. We identified 20 *E. coli* mutants that did not induce *hsp-4::GFP* through the UPR^ER^ reporter (*Pirg-5::GFP*) after three rounds of screening (Table S3). From these 20 *E. coli* mutants, we identified 9 *E. coli* mutants that did not induce *Pirg-5::GFP* through the immunity reporter (*Pirg-5::GFP*) screening (Table S3). (B-C) The bar graph showing that HK-*E. coli* induced *Phsp-4::GFP* (B) and *Pirg-5::GFP* (C) was decreased in animals fed mutant *E. coli* (Heat-killed). Blue numbers are the number of worms scored from at least three independent experiments. Data are represented as mean ± SD. For all panels, * p<0.05, ** p<0.01, *** p < 0.001, **** p<0.0001, ns: no significant difference. Precise P values are provided in Raw Data.

**Figure 3 – figure supplement 2.** Low sugar food, HK-*E. coli*, induce stress response and avoidance behavior in animals. Relative to Figure 3. (A) Venn diagram showing the number of differentially metabolites in HK-*E. coli*-K12, HK-*E. coli*-*yfbR* and *E. coli*. (B) KEGG enrichment analysis of differentially expressed genes in animals fed HK-*E. coli* vs live *E. coli*. We noticed that most of glycolysis/gluconeogenesis genes are up-regulated in animals fed HK-*E. coli*. (C) The bar graph showing that HK-E. coli induced *Phsp-4::GFP* was not affected in animals with D-(+)-trehalose supplementation. Blue numbers are the number of worms scored from at least three independent experiments. Data are represented as mean ± SD. (D) GFP fluorescence images and bar graph showing that HK-*E. coli* induced *Psysm-1::GFP* was decreased in animals with D-(+)-glucose (D-Glc) supplementation. Blue numbers are the number of worms scored from at least three independent experiments. Data are represented as mean ± SD. (E) UPR^ER^ reporter (*Phsp-4::GFP*) expression animals with D-Glc supplementation under OP50, HK-OP50, or NGM condition. (F) immunity reporter (*Pirg-5::GFP*) expression animals with D-Glc supplementation under OP50, HK-OP50, or NGM condition. (G) Development of animals after 48h fed with OP50, HK-OP50, HK-OP50+D-Glc, and HK-OP50+D-GlcA. Blue numbers are the number of worms scored from at least three independent experiments. Data are represented as mean ± SD. For all panels, Scale bar shows on indicated figures, 50 μm. * p<0.05, ** p<0.01, *** p < 0.001, **** p<0.0001, ns: no significant difference. Precise P values are provided in Raw Data.

**Figure 4 – figure supplement 1.** Vitamin C biosynthesis pathway is critical for evaluating low sugar. Relative to Figure 4. (A) The level of total L-ascorbic acid (TAA) in Live-K12, HK-K12, or HK-*yfbR*. Data are represented as mean ±SD from six independent experiments. (B) The level of total L-ascorbic acid (TAA) in animals fed HK-*E. coli* with or without D-Glc supplementation. Data are represented as mean ±SD from eight independent experiments. (C) GFP fluorescence images and bar graph showing that HK-*E. coli* induced *Psysm-1::GFP* was decreased in animals with D-GlcA or vitamin C supplementation. Blue numbers are the number of worms scored from at least three independent experiments. Data are represented as mean ± SD. (D) The bar graph showing that suppression of HK-*E. coli* induced *Psysm-1::GFP* by D-GlcA supplementation was abolished in animals with RNAi of VC biosynthesis genes, which was not affect by vitamin C supplementation. Blue numbers are the number of worms scored from at least three independent experiments. Data are represented as mean ±SD. (E) Food selection assay showing that the preference of HK-*E. coli* with D-GlcA supplementation was abolished in animals by RNAi of vitamin C biosynthesis genes. Data are represented as mean ±SD from six independent experiments, 427-775 animals/assay. For all panels, Scale bar shows on indicated figures, 50 μm. * p<0.05, ** p<0.01, *** p < 0.001, **** p<0.0001, ns: no significant difference. Precise P values are provided in Raw Data.

**Figure 5 – figure supplement 1.** Food behavior of animals. Relative to Figure 5. (A) Food selection assay for OP50 & OP50 + D-Glc, D-GlcA, or VC, respectively. Data are represented as mean ±SD from five independent experiments, 241-1182 animals/assay. (B) Food selection assay for buffer (H_2_O) & buffer (H_2_O) + D-Glc, D-GlcA, or VC, respectively. Data are represented as mean ±SD from five independent experiments, 8-153 animals/assay. (C) Food avoidance assay for N2 & kgb-1 mutant animals fed with HK-*E. coli*. Data are represented as mean ±SD from six independent experiments, 348-660 animals/assay. (D) Translation ability of animals fed with OP50 or HK-OP50 are presented by Western blot of puromycin-labeled peptides. For all panels, * p<0.05, ** p<0.01, *** p < 0.001, **** p<0.0001, ns: no significant difference. Precise P values are provided in Raw Data.

## Supplemental Tables

**Table S1**. Metabolices analysis.

**Table S2**. RNA-seq analysis.

**Table S3**. Screening data for *E. coli* mutant keio library.

**Table S4**. Metabolomics-seq data of HK-K12, HK-*yfbR* and K12.

**Table S5**. RNA-seq data of animals fed with HK-*E. coli* OP50 and *E. coli* OP50.

**Raw-data.** Raw data for experiments.

## Star Methods

### Resource Availability

#### Lead contact

Further information and requests for reagents may be directed to the lead contact Bin Qi.

## Materials availability

All reagents and strains generated by this study are available through request to the lead contact with a completed Material Transfer Agreement.

## Data and code availability

Metabolomics-seq data are accessible in Table S4

RNA-seq data are accessible in Table S5.

This paper does not report original code.

Any additional information required to reanalyze the data reported in this paper is available from the lead contact upon request.

### Experimental model and subject details

#### *C. elegans* strains and maintenance

Nematode stocks were maintained on nematode growth medium (NGM) plates seeded with bacteria (*E. coli OP50*) at 20℃.

1) The following strains/alleles were obtained from the Caenorhabditis Genetics Center (CGC) or as indicated: N2 Bristol (wild type control strain); AU78: *agIs219 [T24B8.5p::GFP::unc-54 3’ UTR + Pttx-3::GFP::unc-54 3’ UTR]*; SJ4005: *zcIs4 [Phsp-4::GFP]*; AY101: *acIs101 [F35E12.5p::GFP + rol-6(su1006)]*; SJ17: *xbp-1 (zc12)*; KU25: *pmk-1(km25)*; AY102: *pmk-1(km25) IV; acEx102 [Pvha-6::pmk-1::GFP + rol-6(su1006)]*; YNU108: *Ex[Prgef-1::pmk-1::GFP;Podr-1::RFP]*^35^; *xbp-1(tm2482)* ^56^; KU21: *kgb-1(km21)*; AU133: *agIs17 [Pmyo-2::mCherry + Pirg-1::GFP] IV*; SJ4100: *zcIs13 [Phsp-6::GFP + lin-15(+)]*.
2) The following strains were constructed by this study: YNU242: *xbp-1(tm2482); pmk-1(km25)* double mutant was constructed by crossing: *xbp-1(tm2482)* with KU25[*pmk-1(km25)*]. YNU240: *ylfEx149 [xbp-1(tm2482); Prgef-1::xbp-1::GFP; Podr-1::RFP]* transgene strain was constructed by injecting plasmid *Prgef-1::xbp-1::GFP* with *Podr-1::RFP* in *xbp-1(tm2482)* background YNU241: *ylfEx150 [xbp-1(tm2482); Pges-1::xbp-1::GFP; Podr-1::RFP]* transgene strain was constructed by injecting plasmid P*ges-1:xbp-1::GFP* with *Podr-1::RFP* in *xbp-1(tm2482)* background

### Bacterial strains

*E. coli-*OP50, *Saprophytic staphylococci*, *Pseudomonas aeruginosa-PA14*, *E. coli-*K12 (BW25113), and *E. coli*-K12 mutant were cultured at 37°C in LB medium. A standard overnight cultured bacteria was then spread onto each Nematode growth media (NGM) plate.

### Culture Medium

MGN Medium: Sigma agar: 20g/L; Bacto Peptone: 2.5g/L; NaCl: 3g/L; MgSO4: 0.12g/L; CaCl: 0.111g/L; PPB: (KH2PO4 0.8M; K2HPO4·3H2O 0.2M) 25ml/L; Cholesterol: 0.005g/L.

LB broth: TPYPTONE: 10g/L; Yeast Extract: 5g/L; NaCl 5g/L.

### Method Details

#### Generation of transgenes

1) To construct the *C. elegans* plasmid for expression of *xbp-1* in neuron, 3057bp promoter of *rgef-1* and genomic DNA of *xbp-1* was inserted into the PPD95.77 vector. DNA plasmid mixture containing *Prgef-1::xbp-1::GFP* (25ng/ul) and *Podr-1::RFP* (25ng/ul) was injected into the gonads of adult *xbp-1(tm2482)*.
2) To construct the *C. elegans* plasmid for expression of *xbp-1* in intestine, 2549 bp promoter of ges-1 and genomic DNA of *xbp-1* was inserted into the PPD95.77 vector. DNA plasmid mixture containing *Pges-1::xbp-1::GFP* (25ng/ul) and *Podr-1::RFP (25ng/ul)* was injected into the gonads of adult *xbp-1(tm2482)*.

### Preparation and feeding of worm food

We followed an established protocol ^24, 25^ to prepare heat-killed (HK) *E. coli*. Briefly, a standard OD_600_=0.5-0.6 of *E. coli* OP50 and *E. coli* K12 grown in LB broth was concentrated to 1/20 vol and was then heat-killed at 80℃ for 180 min. About 150ul of the heat-killed bacteria was spread onto each 35mm NGM plate.

### Preparation of HK-*E. coli* + carbohydrate or vitamin C food

1) 100ul of water, 100ul of L-ascorbic acid (dissolved in water at a concentration of 100mg/ml, Sangon Biotech, 100143-0100) or 100 ul of D-glucuronic acid (dissolved in water at a concentration of 100mg/ml, Adamas, 1102520) was mixed with 500 ul of HK-*E. coli*, then 150ul of the mixture was spread onto 35mm NGM plates.
2) 12.5ul of water or 12.5ul of D-(+)-glucose (dissolved in water at a concentration of 100mg/ml, Sangon Biotech, A501991-0500) was mixed with 500 ul of HK-*E. coli*, then 150ul of the mixture was spread onto 35mm NGM plates.

### Behavioral assay

#### *C. elegans* selection assays

For *C. elegans* to have enough space to evaluate food, we add 18ul of the sample onto a 35mm NGM plate. This creates a round lawn with a radius of 5mm, which occupies about 8% of the total plate area.

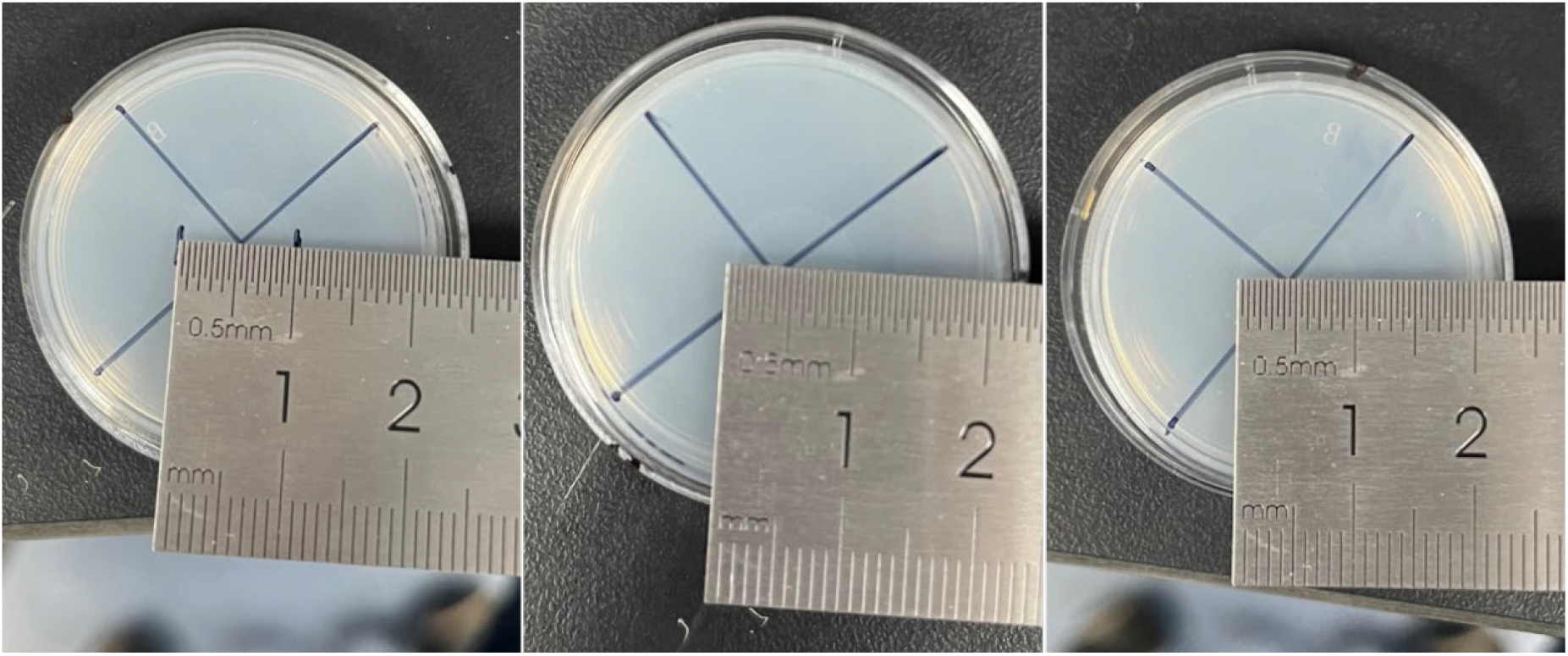

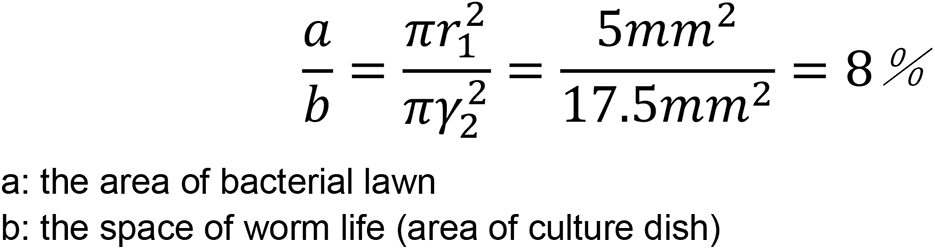

1) 18ul of heat-killed OP50, live OP50, and LB broth (as the buffer for bacteria) was added into 35mm NGM plate in an equilateral triangle pattern. Then, synchronized L1 worms were seeded in the center of NGM plate for 16-17h at 20℃ (as indicated in Figure1B).
2) 18ul of heat-killed OP50 and heat-killed OP50 with D-GlcA or vitamin C was added into 35mm NGM plate in an equilateral triangle pattern, then synchronized L1 worms were seeded on equilateral triangle of NGM plate for 16-17h at 20℃ (as indicated in Figure 5A).

Here is Selection rate formula:

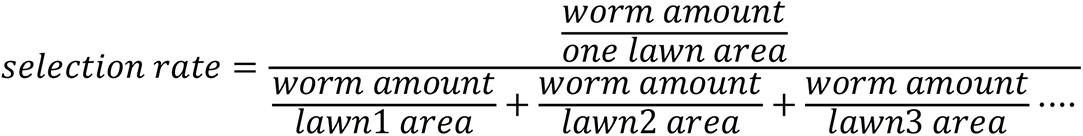

#### *C. elegans* aversion assays

18ul food was spread out the center of NGM plate, then synchronized L1 by bleach solution (NaOH: 1M, NaClO:4-6%) worms were seeded on center of food for 16-17h at 20 ℃.

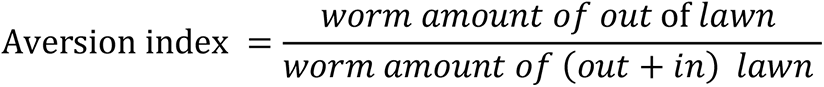

Three to ten replicates for each condition were performed for each assay, and the experiments were duplicated on different days.

### Analysis of the fluorescence intensity in worms

The synchronized L1 worms carrying either UPR^ER^ reporter (*Phsp-4::GFP*) or innate immunity reporter (*Pirg-5::GFP*; *Psysm-1p::GFP*; *Pirg-1p::GFP*) were seeded on NGM with indicated food and incubated for 24h at 20℃. For fluorescence imaging, worms were anesthetized with 25 mM levamisole and photographed using either an Olympus BX53 microscope or Olympus MVX10 dissecting microscope equipped with a DP80 camera.

The fluorescence intensity in entire intestinal region was quantified using ImageJ software and normalized to the body area.

#### *E. coli* Keio collection screen

The whole Keio *E. coli* single mutant collection (Baba et al., 2006) was screened. Mutant bacteria strains, as well as the wild-type control strain BW25113, were cultured in LB medium with 50µg/ml kanamycin at 37℃ until an OD_600_ of 0.5-0.6 was reached. The bacteria were then heat-killed following our established protocol ^24^, and 150 ul of the heat-killed mutant *E. coli* was spread onto 35 mm NGM plates. Synchronized L1 worms carrying UPR^ER^ reporter (*Phsp-4::GFP*) were seeded and cultured for 24h at 20℃. The fluorescence was then examined by using an Olympus MVX10 dissecting microscope, progressive screening three times. Next, a 4^th^ screen was performed using immune reporter (*Pirg-5::GFP*) animals fed with HK-E. coli mutants that reduced the *Phsp-4::GFP* fluorescence. Finally, an aversion behavior assay was performed using HK-*E. coli* mutants that both reduced *Phsp-4::GFP* and *Pirg-5::GFP.* HK-*E. coli* mutants that reduced UPR^ER^, immune and avoidance behavior were identified through this screening.

### RNAi treatment

RNAi plasmid is delivered in a *E. coli* strain, HT115, from either the MRC RNAi library ^57^ or the ORF-RNAi Library ^58^. RNAi plates were prepared by adding IPTG to NGM agar to a final concentration of 1 mM. Overnight *E. coli* cultures (LB broth containing 100 ug/ml ampicillin and 100uM IPTG) of specific RNAi strains and the control HT115 strain were seeded onto RNAi feeding plates and cultured at room temperature until dry. Synchronized L1 worms were treated RNAi by feeding (Ahringer, Reverse genetics, WormBook 2006) for the first generation and allowed to grow to maturity. The worms were then bleached and hatched in M9 buffer for 18hr. The synchronized L1 worms were then seeded on the indicated feeding plate.

### Western blot

To measure the level of p-PMK-1, worms (feeding different food for 4h) were analyzed by standard western blot methods and probed with anti-p38 (dilution = 1:5,000; Cell Signaling, 9212S), anti-p-p38 (dilution = 1:5,000; Cell Signaling, 4511S) and anti-α-tubulin (dilution = 1:10,000; Sigma T5168) as a loading control.

To measure the level of protein translation, worms (feeding different food for 24h) were analyzed by standard western blot methods and probed with anti-Puromycin (dilution = 1:10,000; Sigma-Aldrich, MABE343) and anti-α-tubulin (dilution = 1:10,000; Sigma T5168) as a loading control.

### Total content of ascorbic acid (TAA) assay

The total content of ascorbic acid was measured using the kits (Beijing Biotech-Pack-analytical Scientific Co., Ltd., Beijing, China, BKWB132 http://biotech-pack-analytical.foodmate.net/) according to the manufacturer’s protocol. Briefly, L1 worms were seeded on the different feeding assay plate and cultured for 4 hours. The worms were then lysed in ice-cold conditions using lysis buffer. Equal amounts of protein were used for the normalization. Here is formula for getting TAA concentration

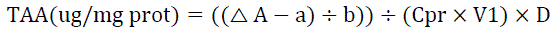

V1 – the volume of supernatant of for experiment

Cpr – the concentration of supernatant protein

D – Dilution ratio of supernatant

a – the intercept of standard curve

b – the slope of standard curve

the standard curve y = 0.0611x + 0.0003 for Figure 4C; y = 0.0258x + 0.0066 for Figure 4B, Figure 4-figure supplement 1A and Figure 4-figure supplement 1B.

### Preparation of samples for RNA sequencing

RNA-seq was done with three biological replicates that were independently generated, collected, and processed. Adult wild type (N2) worms were bleached and then the eggs were incubated in M9 for 18 hours to obtain synchronized L1 worms. L1 worms were cultured in the NGM plate with *HK-E. coli* or *E. coli* for 4hrs at 20℃. L1 worms were then collected for sequencing.

### RNA sequencing and data processing

For the RNA sequencing assay, cDNA libraries were constructed, and single-end libraries were sequenced using the Illumina platform (Novogene, Beijing, China). HISAT2 ^59^ was used to map the clean reads to the reference gene sequence (Species: Caenorhabditis_elegans; Source: NCBI; Reference Genome Version: GCF_000002985.6_WBcel235), and then “featureCounts” tool in subread software ^60^ was used to calculate the gene expression level of each sample. Read counts were inputted into DESeq2 ^61^to calculate differential gene expression and statistical significance. Differentially expressed genes (DEGs) were screened using following criteria: |log2(FoldChange)| > 1 & padj<= 0.05.

### Preparation of samples for metabolome sequencing

Metabolome-seq of bacterial was done with four biological replicates that were independently generated, collected, and processed. Total of three group E. coli sample including: *E. coli K12* (Con), HK-*E. coli K12* (HK), and HK-*E*. *coli yfbR* mutant (HY). All bacteria were overnight cultured to the same OD (OD_600_=1). *E. coli K12* and *E. coli yfbR* mutant are heat-killed (80, 180min), *E. coli* K12, HK-*E. coli* K12 and HK-*E. coli yfbR* mutant were then spread out NGM plate for 72hs at room temperature. Finally, sample was collected into 1.5ml tube by using sterile cell scraping.

### Metabolome sequencing and data processing

Metabolome were sequenced using the Ultra Performance Liquid Chromatography (UPLC) (ExionLC AD, https://sciex.com.cn/) and Quadrupole-Time of Flight (TripleTOF 6600, AB SCIEX) for Non-targeted; Ultra Performance Liquid Chromatography (UPLC) (ExionLC AD, https://sciex.com.cn/) and Tandem mass spectrometry (MS/MS) (QTRAP®, https://sciex.com/) for Broad targeting (Metware, Wuhan, China). Multiple reaction monitoring (MRM) was used to calculate the expression level of each metabolite. Differential metabolites were screened through Fold change ≥ 2 or Fold change ≤ 0.5 and VIP ≥ 1 (Variable Importance in Projection of OPLS-DA model).

### Microscopy

Analysis of fluorescence was performed with an Olympus BX53 microscope, CLSM (Zeiss LSM900), or Olympus MVX10 dissecting with a DP80 camera.

## QUANTIFICATION AND STATISTICAL ANALYSIS

### Quantification

ImageJ software was used for quantifying fluorescence intensity of UPR^ER^ and Innate immunity reporter. ImageJ software was used for counting the number of worms about selection and aversion behavior.

### Statistical analysis

All statistical analyses were performed in Graphpad prism 8.0. Two-tailed unpaired t test was used for statistical analysis of two groups of samples, one-way or two-way ANOVA was used for statistical analysis of more than two groups of samples. Data are presented as Mean ± SD, and p<0.05 was considered a significant difference, “*” represents p<0.05, “**” represents p<0.01, “***” is represents < 0.001, “****” represents p<0.0001, “ns” represents no significant difference. For all figures, ‘‘n’’ represents the number of worms scored from at least three independent experiments.

## Notes

### Competing Interest Statement

The authors have declared no competing interest.

### Summary of Updates

In this revision, we added to the discussion about how is the activation of the PMK-1 pathway driven by/coordinated with UPR activation？ We completed the control group of section data. (Figure 3C-F) We tested animals development in HK-E.coli, HK-E. coli+D-Glc/D-GlcA (figure 3-figure supplement 2G)

