## Supplemental Figure for "UPR^ER^–immunity axis acts as physiological food evaluation system that promotes aversion behavior in sensing low-quality food"

Figure 1-figure supplement 1

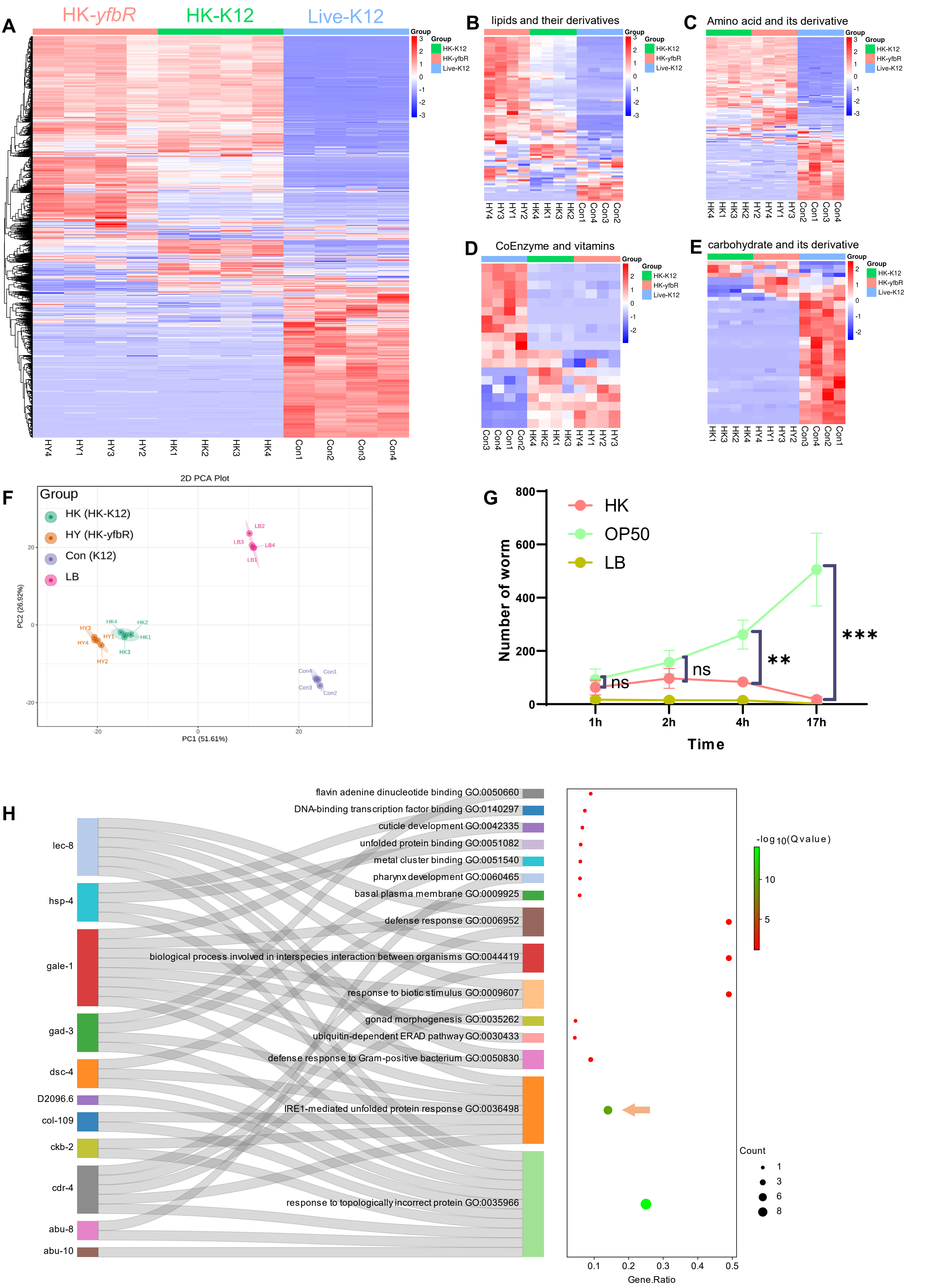

Figure 1-figure supplement 2

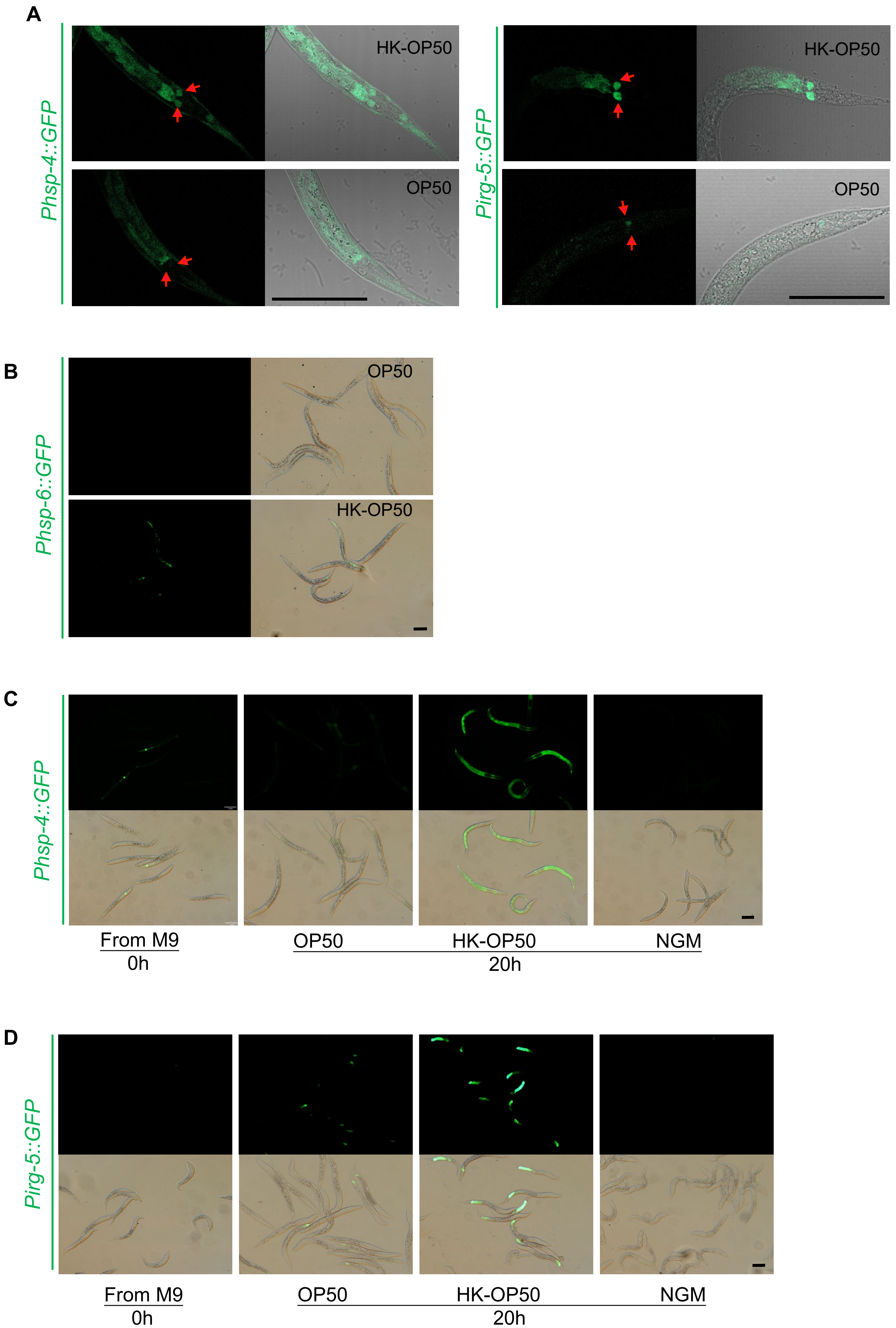

**Figure 1 – figure supplement 1. Food selection assay of animals fed HK-*E. coli* or *E. coli.* Relative to Figure 1.**

(A-E) Metabolomics analysis of different quality food (HK-K12, HK-*yfbR* and Live-K12). Cluster analysis of all metabolites (A), lipids and their derivatives (B), amino acids and their derivatives (C), coenzymes and vitamins (D), and carbohydrates and their derivatives (E) from Live-K12, HK-K12, and HK-*yfbR*. Color indicates the relative level of each metabolite. HK-K12: heat-killed *E. coli* wild-type K12; HK-*yfbR:* heat-killed *E. coli mutant yfbR;* K12: live *E. coli* wild-type K12*.* z-score for standardizing data, complete for bi-clustering algorithm, and Euclidean for distance method.

(F) Principal component analysis to test the repeatability of the metabolic experiment. HK-K12: heat-killed *E. coli* wild-type K12; HK-*yfbR*: heat-killed *E. coli* mutant *yfbR*; K12: live *E. coli* wild-type K12. LB: LB buffer for culturing *E. coli.*

(G) The number of worms in each position calculated at the indicated time, indicating that animals initially select both foods (1-2h), but eventually favor high-quality food (Live E. coli) until 17h.

(H) GO enrichment analysis of $UPR^{ER}$ dependent IRE-1 branch.

For all panels, * p<0.05, ** p<0.01, *** p < 0.001, **** p<0.0001, ns: no significant difference. Precise P values are provided in Raw Data.

**Figure 1 – figure supplement 2. Stress response of animals fed HK-*E. coli* or *E. coli.* Relative to Figure 1.**

(A) $UPR^{ER}$ reporter (*Phsp-4::GFP*) and immunity reporter (*Pirg-5::GFP*) were induced in the neuron and intestine of animals fed with HK-*E. coli.* Neurons are highlighted with a red arrow.

(B) $UPR^{mit}$ reporter (*Phsp-6::GFP*) was weakly induced in animals fed with HK- *E. coli.*

(C) $UPR^{ER}$ reporter (*Phsp-4::GFP*) expression in animals under normal food (OP50), low quality food (HK-OP50), and starved (M9: hatching L1 worm in M9, NGM) condition, L1 animals were cultured in OP50, HK-OP50 or starved NGM for 20h.

(D) Immunity reporter (*Pirg-5::GFP*) expression in animals under normal food (OP50), low quality food (HK-OP50), and starved (M9: hatching L1 worm in M9, NGM), L1 animals were cultured in OP50, HK-OP50 or starved NGM for 20h.

For all panels, Scale bar shows on indicated figures, 50 μm.

### Figure 2-figure supplement 1

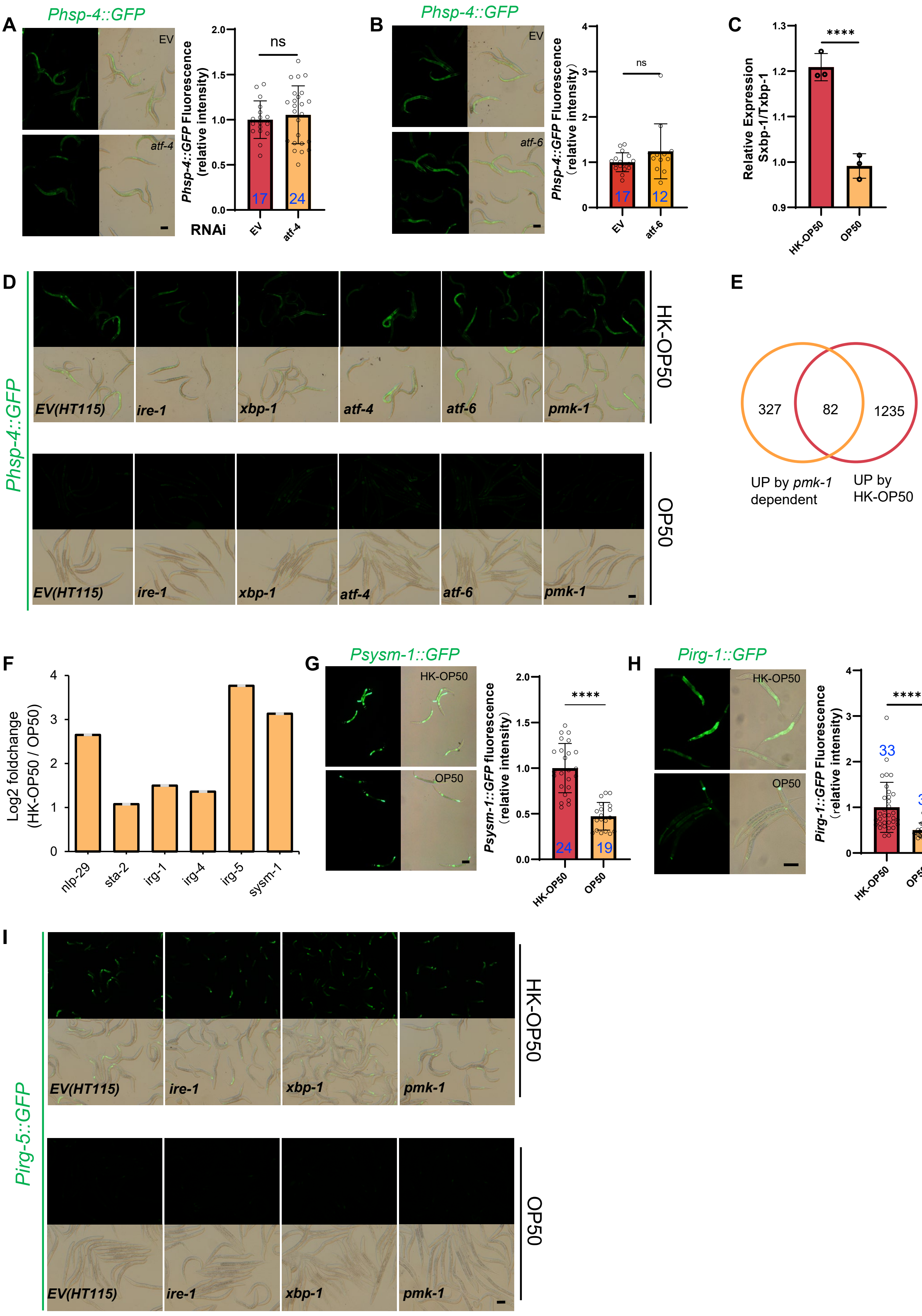

### Figure 2-figure supplement 2

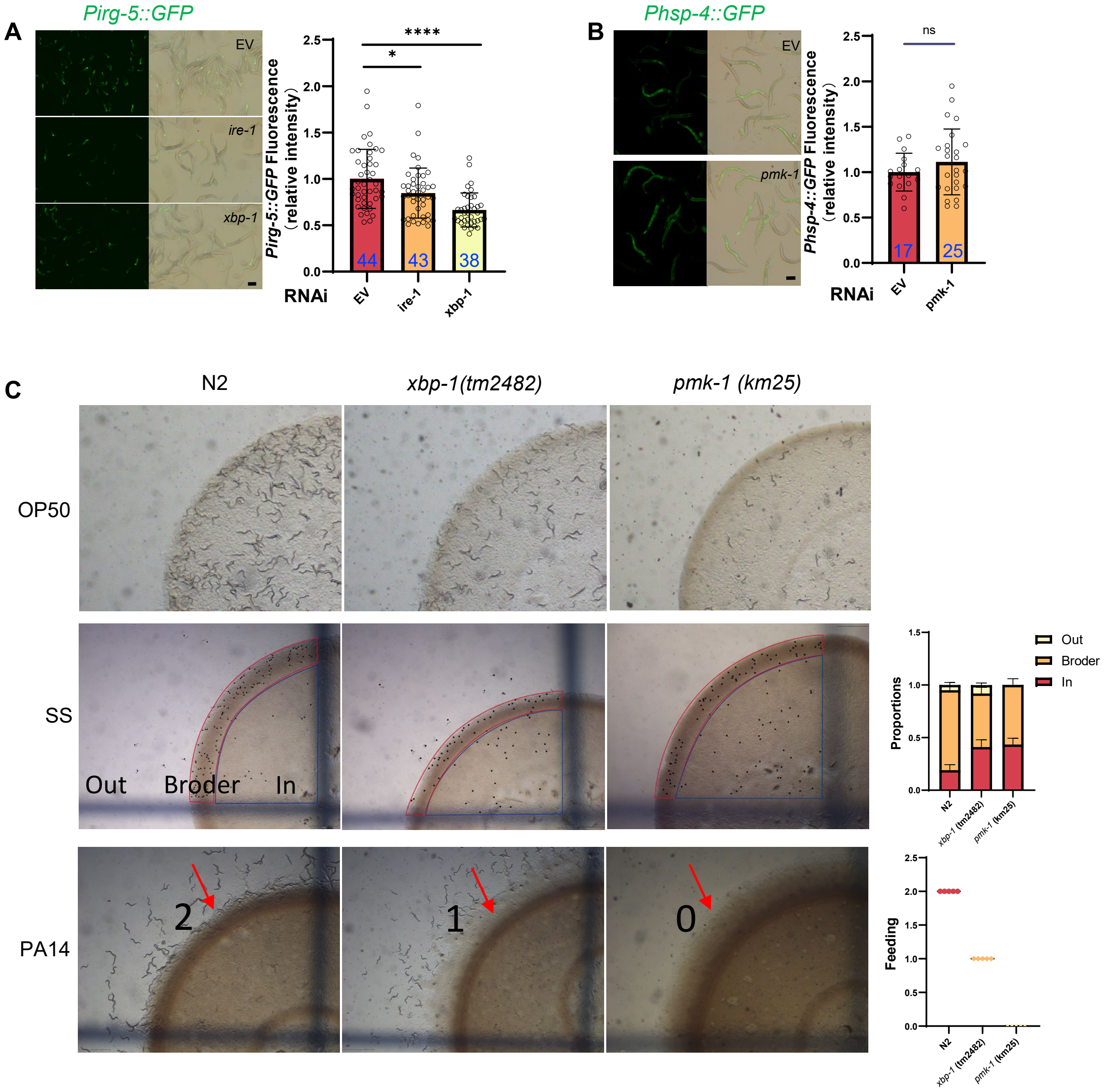

**Figure 2 - figure supplement 1. UPR$^{ER}$ and innate immunity pathway in animals are critical for evaluating HK-*E. coli*. Relative to Figure 2.**

(A-B) GFP fluorescence images and bar graph showing that HK-*E. coli* induced *Phsp-4::GFP* was not affected in animals with *atf-4* (A) or *atf-6* (B) RNAi treatment. Blue numbers are the number of worms scored from at least three independent experiments. Data are represented as mean ± SD.

(C) qPCR showing that IRE-1-mediated splicing of *xbp-1* mRNA is induced in animals fed with HK-*E. coli.*

(D) UPR$^{ER}$ reporter (*Phsp-4::GFP*) expression in animals with candidate RNAi feeding OP50 or HK-OP50.

(E) Venn diagram showing the numbers of PMK-1 dependent genes [30] and up-regulated genes in animals fed HK-*E. coli,* and their overlap.

(F) The expression of PMK-1 dependent genes which was extracted from RNA-seq data from animals fed with HK-*E. coli*. The data from average of three independent experiments.

(G-H) GFP fluorescence images and bar graph showing that *Psysm-1::GFP* (G) and *Pirg-1::GFP* (H) were induced in animals fed HK-*E. coli.* Blue numbers are the number of worms scored from at least three independent experiments. Data are represented as mean ± SD.

(I) Immunity reporter (*Pirg-5::GFP*) expression in animals with candidate RNAi feeding OP50 or HK-OP50.

For all panels, Scale bar shows on indicated figures, 50 μm. * p<0.05, ** p<0.01, *** p < 0.001, **** p<0.0001, ns: no significant difference. Precise P values are provided in Raw Data.

**Figure 2 - figure supplement 2. UPR$^{ER}$ positively regulates innate immunity pathway in animals. Relative to Figure 2.**

(A) GFP fluorescence images and bar graph showing that HK-*E. coli* induced *Pirg-5::GFP* was decreased in animals with *ire-1* or *xbp-1* RNAi treatment. Blue numbers the number of worms scored from at least three independent experiments. Data are represented as mean ± SD.

(B) GFP fluorescence images and bar graph showing that *Phsp-4::GFP* was not affected in animals with *pmk-1* RNAi treatment. Blue numbers the number of worms scored from at least three independent experiments. Data are represented as mean ± SD.

(C) Images and bar graph showing that the avoidance behavior of N2, *xbp-1* mutant, and *pmk-1* mutant in diverse food (*Saprophytic staphylococci* - SS, *Pseudomonas aeruginosa* - PA14, or OP50).

"Broder" indicates regions with thicker SS boundaries; "In" denotes areas inside SS ; "Out" refers to areas without SS. The scale is as follows: 2 for animals capable of escaping from PA14 and feeding on PA14 located with edges; 1 for animals capable of escaping from PA14 but not feeding on PA14 located with edges; 0 for animals unable to escape from PA14 and being killed by PA14. Data are represented as mean ±SD from five independent experiments, 359-670 animals/assay.

For all panels, Scale bar shows on indicated figures, 50 μm. * p<0.05, ** p<0.01, *** p < 0.001, **** p<0.0001, ns: no significant difference. Precise P values are provided in Raw Data.

Figure 3-figure supplement 1

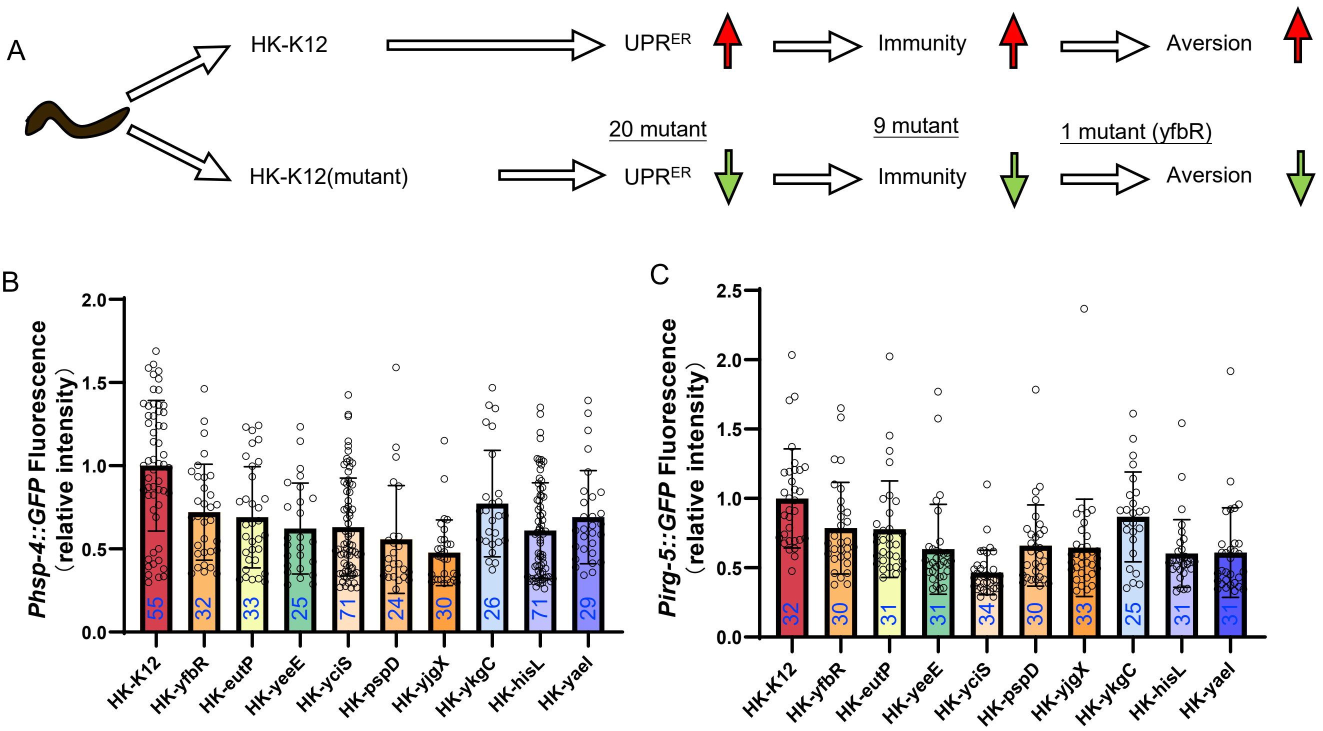


**Figure 3 - figure supplement 1. *E. coli* Keio mutant screening. Relative to Figure 3.**

(A) Flow chart of strategy for *E. coli* Keio mutant screening. We identified 20 *E. coli* mutants that did not induce *hsp-4::GFP* through the UPR$^{ER}$ reporter (*Pirg-5::GFP*) after three rounds of screening (Table S3). From these 20 *E. coli* mutants, we identified 9 *E. coli* mutants that did not induce *Pirg-5::GFP* through the immunity reporter (*Pirg-5::GFP*) screening (Table S3).

(B-C) The bar graph showing that HK-*E. coli* induced *Phsp-4::GFP* (B) and *Pirg-5::GFP* (C) was decreased in animals fed mutant *E. coli* (Heat-killed). Blue numbers the number of worms scored from at least three independent experiments. Data are represented as mean ± SD.

For all panels, * p<0.05, ** p<0.01, *** p < 0.001, **** p<0.0001, ns: no significant difference. Precise P values are provided in Raw Data.

### Figure 3-figure supplement 2

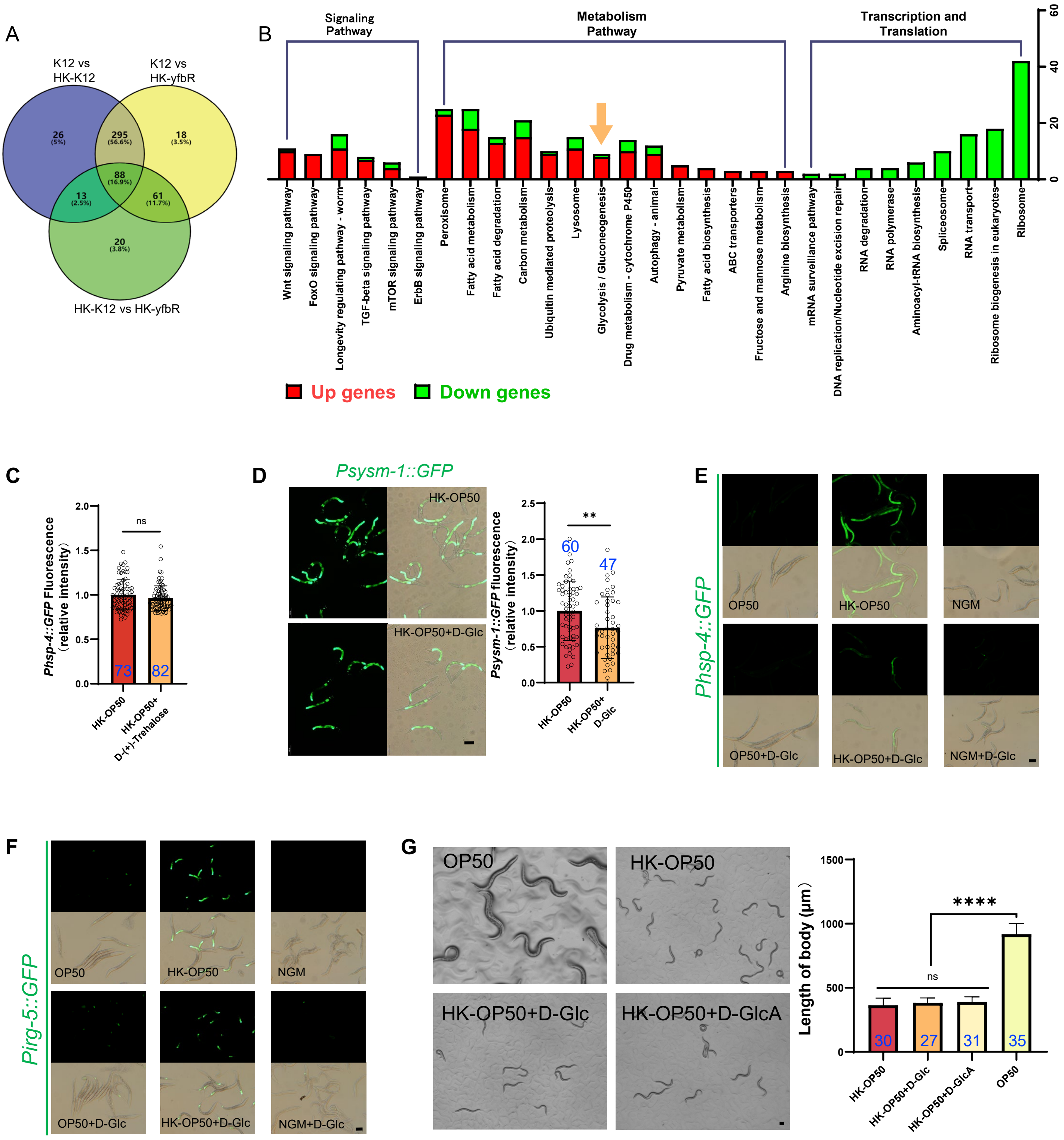

**Figure 3 - figure supplement 2. Low sugar food, HK-*E. coli*, induce stress response and avoidance behavior in animals. Relative to Figure 3.**

(A) Venn diagram showing the number of differentially metabolites in HK-*E. coli*-K12, HK-*E. coli*-*yfbR* and *E. coli*.

(B) KEGG enrichment analysis of differentially expressed genes in animals fed HK-*E. coli* vs live *E. coli*. We noticed that most of glycolysis/gluconeogenesis genes are up-regulated in animals fed HK-*E. coli*.

(C) The bar graph showing that HK-E. coli induced *Phsp-4::GFP* was not affected in animals with D-(+)-trehalose supplementation. Blue numbers the number of worms scored from at least three independent experiments. Data are represented as mean ± SD.

(D) GFP fluorescence images and bar graph showing that HK-*E. coli* induced *Psysm-1::GFP* was decreased in animals with D-(+)-glucose (D-Glc) supplementation. Blue numbers the number of worms scored from at least three independent experiments. Data are represented as mean ± SD.

(E) $UPR^{ER}$ reporter (*Phsp-4::GFP*) expression animals with D-Glc supplementation under OP50, HK-OP50, or NGM condition.

(F) immunity reporter (*Pirg-5::GFP*) expression animals with D-Glc supplementation under OP50, HK-OP50, or NGM condition.

(G) Development of animals after 48h fed with OP50, HK-OP50, HK-OP50+D-Glc, and HK-OP50+D-GlcA. Blue numbers the number of worms scored from at least three independent experiments. Data are represented as mean ± SD.

For all panels, Scale bar shows on indicated figures, 50 μm. * p<0.05, ** p<0.01, *** p < 0.001, **** p<0.0001, ns: no significant difference. Precise P values are provided in Raw Data.

### Figure 4-figure supplement 1

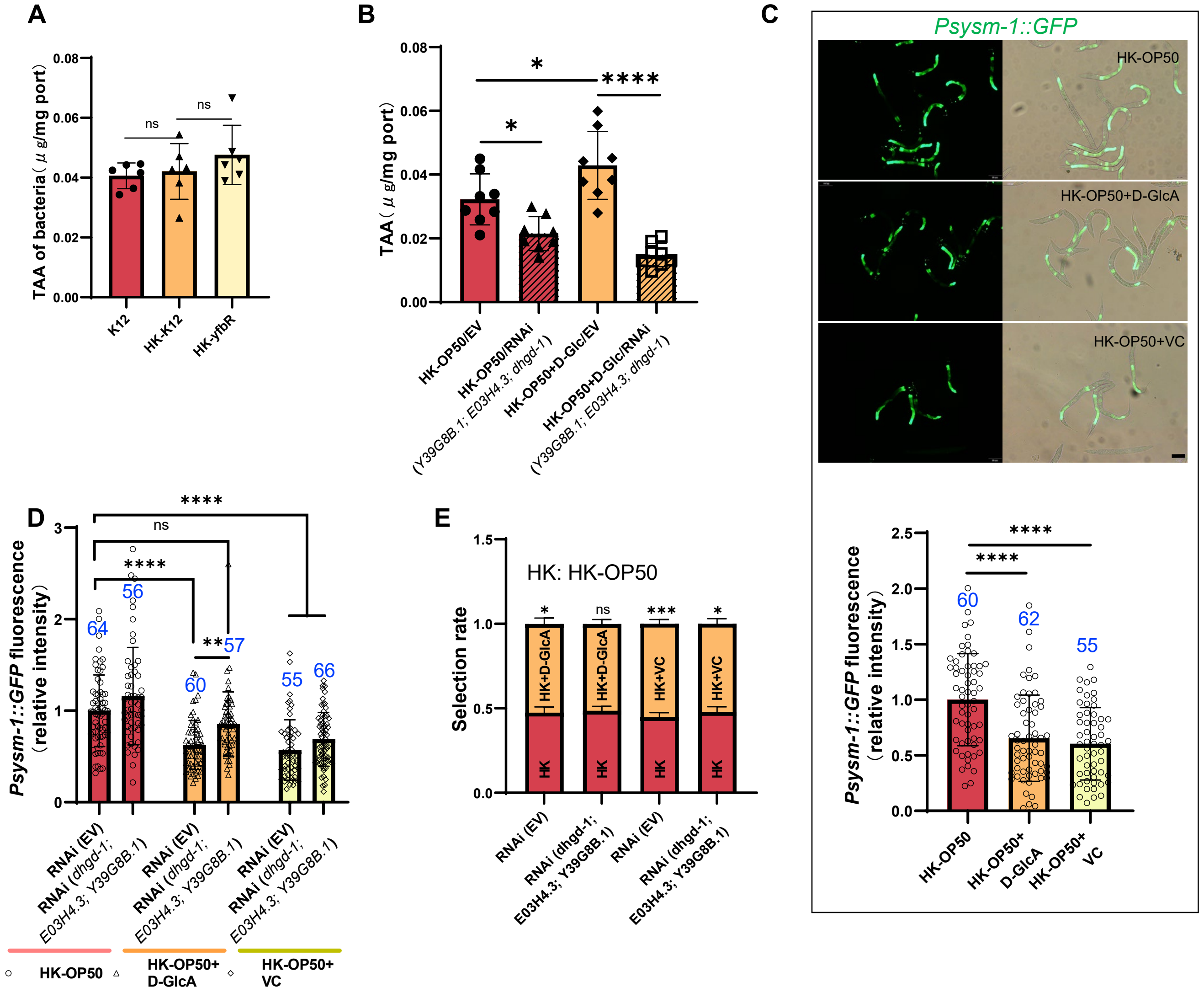

**Figure 4 - figure supplement 1. Vitamin C biosynthesis pathway is critical for evaluating low sugar. Relative to Figure 4.**

(A) The level of total L-ascorbic acid (TAA) in Live-K12, HK-K12, or HK-*yfbR*. Data are represented as mean ±SD from six independent experiments.

(B) The level of total L-ascorbic acid (TAA) in animals fed HK-*E. coli* with or without D-Glc supplementation. Data are represented as mean ±SD from eight independent experiments.

(C) GFP fluorescence images and bar graph showing that HK-*E. coli* induced *Psysm-1::GFP* was decreased in animals with D-GlcA or vitamin C supplementation. Blue numbers the number of worms scored from at least three independent experiments. Data are represented as mean ± SD.

(D) The bar graph showing that suppression of HK-*E. coli* induced *Psysm-1::GFP* by D-GlcA supplementation was abolished in animals with RNAi of VC biosynthesis genes, which was not affect by vitamin C supplementation. Blue numbers the number of worms scored from at least three independent experiments. Data are represented as mean ±SD.

(E) Food selection assay showing that the preference of HK-*E. coli* with D-GlcA supplementation was abolished in animals by RNAi of vitamin C biosynthesis genes. Data are represented as mean ±SD from six independent experiments, 427-775 animals/assay.

For all panels, Scale bar shows on indicated figures, 50 μm. * p<0.05, ** p<0.01, *** p < 0.001, **** p<0.0001, ns: no significant difference. Precise P values are provided in Raw Data.

Figure 5-figure supplement 1

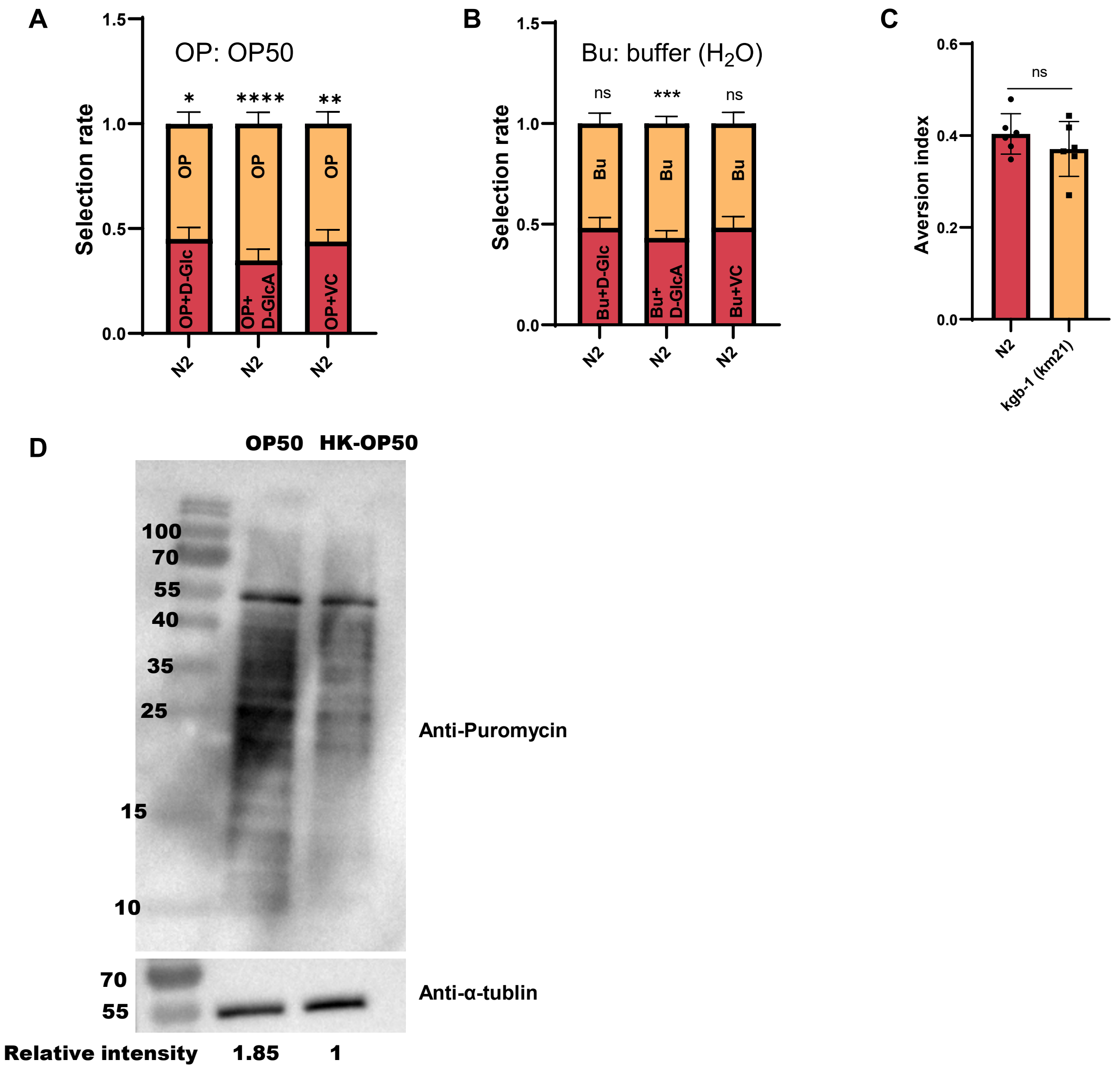

**Figure 5 - figure supplement 1. Food behavior of animals. Relative to Figure 5.**

(A) Food selection assay for OP50 & OP50 + D-Glc, D-GlcA, or VC, respectively. Data are represented as mean ±SD from five independent experiments, 241-1182 animals/assay.
(B) Food selection assay for buffer ($H_2O$) & buffer ($H_2O$) + D-Glc, D-GlcA, or VC, respectively. Data are represented as mean ±SD from five independent experiments, 8-153 animals/assay.
(C) Food avoidance assay for N2 & kgb-1 mutant animals fed with HK-*E. coli*. Data are represented as mean ±SD from six independent experiments, 348-660 animals/assay.
(D) Translation ability of animals fed with OP50 or HK-OP50 are presented by Western blot of puromycin-labeled peptides.

For all panels, * p<0.05, ** p<0.01, *** p < 0.001, **** p<0.0001, ns: no significant difference. Precise P values are provided in Raw Data.
